# HuR modulation with tanshinone mimics impairs LPS response in murine macrophages

**DOI:** 10.1101/2023.01.16.524289

**Authors:** Isabelle Bonomo, Giulia Assoni, Valeria La Pietra, Giulia Canarutto, Elisa Facen, Greta Donati, Chiara Zucal, Silvia Genovese, Mariachiara Micaelli, Anna Pérez-Ràfols, Sergio Robbiati, Dimitris L. Kontoyannis, Marilenia De Matteo, Marco Fragai, Pierfausto Seneci, Luciana Marinelli, Daniela Arosio, Silvano Piazza, Alessandro Provenzani

## Abstract

Lipopolysaccharide exposure to macrophages induces an inflammatory response that is heavily regulated at the transcriptional and post-transcriptional levels. HuR (ELAVL1) is an RNA binding protein that binds and regulates the maturation and half-life of AU/U rich elements (ARE) containing cytokines and chemokines transcripts, mediating the LPS-induced response. Here we investigated how and to what extent small molecule tanshinone mimics (TMs) inhibiting HuR-RNA interaction counteract LPS stimulus in macrophages. We show TMs exist in solution in keto-enolic tautomerism and that, by molecular dynamic calculations, the orto quinone form is the bioactive species interacting with HuR and inhibiting its binding mode vs mRNA targets. A chemical blockage of the diphenolic, reduced form as a diacetate caused the loss of activity of TMs *in vitro* but resulted to prodrug-like activity *in vivo*. The murine macrophage cell line RAW264.7 was treated with LPS and TMs, and the modulation of cellular LPS-induced response was monitored by RNA and Ribonucleoprotein immunoprecipitation sequencing. Correlation analyses indicated that LPS induced a strong coupling between differentially expressed genes and HuR-bound genes, and that TMs reduced such interactions. Functional annotation addressed a specific set of genes involved in chemotaxis and immune response, such as *Cxcl10, Il1b, Cd40*, and *Fas*, with a decreased association with HuR, a reduction of their expression and protein secretion. The same effect was observed in primary murine bone marrow-derived macrophages, and *in vivo* in an LPS induced peritonitis model, in which the serum level of Cxcl10 and Il1b was strongly reduced, endowing TMs such as **TM7nox** with remarkable anti-inflammatory properties *in vivo*.

## Introduction

RNA Binding Proteins (RBPs) play a pivotal role in the regulation of gene expression in eukaryotes by exploiting RNA-protein and protein-protein interactions^1,2^. RBPs’ aberrant expression, their modulation or mis-localization lead to the insurgence of complex phenotypes and diseases^3–5^. Therefore, targeting and modulating the activity of RBPs associated to various pathologies represents a new promising therapeutic strategy^6^. In this context, the Human antigen R (HuR, official name ELAVL1) is among the most widely studied RBPs. It belongs to the ELAVL protein family, is ubiquitously expressed in human tissues and highly conserved during mammalian evolution^7–9^. HuR binds AU/U rich elements (AREs), located mainly in the 3’UTRs of coding and non-coding RNA. ARE sequences are found in 7% of the human mRNAs, coding for proteins involved in key cellular processes as immune response and inflammation^10–13^, cell division and proliferation^14,15^, angiogenesis^16–18^, senescence^14,19,20^ and apoptosis^15,21,22^. A strong regulatory role for HuR is proven by the fact that ≈90% of mRNAs coding for cytokines and chemokines contain repeated AREs sites in their 3’UTR^23–25^. Consequently, HuR aberrant expression or sub-cellular distribution is connected to diseases such as cancer and immune pathologies^26,27^. Thus, HuR should represent a valuable therapeutic target. HuR structure is characterized by three RNA recognition motifs (RRMs). RRM1 and RRM2 strictly cooperate, with RRM1 primarily responsible for RNA binding, while RRM2 and the inter-domain linker significantly increase the RNA binding affinity of both RRMs. In presence of RNA, RRM1 and RRM2 undergo conformational changes, assume a closed shape and form a positively charged cleft responsible for RNA binding^28^. RRM3 is known to bind mRNA polyA tails and promote protein oligomerization, but also contributes to RNA binding^29^.

Among HuR inhibitors reported so far^9^, Tanshinone Mimics (TMs) are synthetic compounds interfering with HuR activity, whose Structure Activity Relationships (SARs) were described^30^. They modulate HuR activity by competing with HuR:RNA complex formation, interacting with the protein in the region responsible for RNA binding^30,31^. Consequently, TMs inhibit HuR regulation of cancerogenic mRNAs, showing anti-tumoral traits. Here, we explore TMs’ inhibitory activity on HuR-mediated response to inflammatory stimuli, specifically driven by LPS. HuR up-regulates inflammation processes through several mechanisms, such as impeding toll-like receptor 4 (TRL4) mRNA degradation and stabilizing various inducible pro-inflammatory transcripts (e.g interferon γ, TNFα, IL-6)^32^. PAR-CLIP (PhotoActivatable Ribonucleoside-enhanced CrossLinking and ImmunoPrecipitation) experiments in primary macrophages challenged with an LPS inflammatory stimulus showed the existence of a complex post-transcriptional response driven by the engagement of the RBP Tristetraprolin (TTP) and HuR^33^. Here, we synthesized three TM derivatives and a diacetate prodrug to improve their solubility and provide significant *in vivo* activity. The mode of action of such TMs in counteracting the LPS response during co-administration was elucidated at the genome-wide level, and led to the identification of inflammatory target mRNAs, as *Cxcl10* and *Il1b*, whose HuR binding is regulated by TMs. Moreover, the anti-inflammatory properties of TMs are shown *in vivo*, in an LPS-induced peritonitis mouse model.

## Materials and Methods

### Chemical Synthesis

Synthetic procedures and compounds characterization are reported in the Supplementary Material.

### Homogeneous Time Resolved Fluorescence (HTRF)

A recombinant M1M2 version of HuR has been expressed in *E*.*coli BL21* cells, as previously described^31,34,35^. Homogeneous Time Resolved Fluorescence was used to determine the inhibition of the binding between His-tag M1M2 HuR recombinant protein and a 26 biotinylated TNF ARE RNA probe (5’-AUUAUUUAUUAUUUAUUUAUUAUUUA-3’) by TMs. We calculated the protein Hook point, by testing different concentrations of the protein (0, 5 nM, 10 nM, 20 nM, 50 nM, 100 nM, 200 nM and 500 nM) in presence of an ARE probe (50 nM). After inhibition, tests were performed with the HuR construct (20 nM) and the probe (50 nM), HTRF experiments were performed according to the manufacturer instructions (Cisbio). Signals were measured with a Tecan Spark microplate reader, using the protocol indicated by the manufacturer (Cisbio).

### RNA-Electrophoretic Mobility Shift Assays (REMSAs)

M1M2 HuR recombinant protein was purified and REMSAs were performed as previously described^31,34,35^. Briefly, the protein (3.7 nM) was incubated for 30 min with 5′-DY681-labeled AU-rich RNA probe (1 nM) and DMSO as control, or with TMs at various dosages. Afterwards, samples were loaded on 4% native polyacrylamide gel; images were developed with a Typhoon Trio scanner (GE Healthcare) at the resolution for the DY681-probe.

### Cell line primary cultures

Murine macrophage RAW264.7 (ICLC, Genova) cell lines were maintained in high glucose Dulbecco’s modified Eagle’s medium (DMEM) by adding 10% fetal bovine serum (FBS, Lonza), 2 mM L-glutamine, 100 U/mL penicillin-streptomycin (Lonza) in standard growth conditions. Murine bone marrow–derived macrophages (BMDMs) were obtained from sex balanced C57BL6/j 6-12 weeks old mice according to published protocols^36,37^. To stimulate differentiation, BMDMs were cultured for 7 days with 10% of supernatant medium from L929 fibroblasts (Sigma 85011425), and maintained in Roswell Park Memorial Institute (RPMI) 1640 Medium by adding 5% fetal bovine serum (FBS, Lonza), 2 mM L-glutamine, and 100 U/mL penicillin-streptomycin (Lonza) in standard growth conditions^36,37^.

### Animal inflammation models and sera collection

C57BL6/j mice were purchased from Charles River Laboratories, bred and maintained in the animal facilities of the Department CIBIO under pathogen-free conditions and according to the authorization received from the Italian Health Ministry ethical committee for animal experimentation #629-2018. To measure inflammatory factors secretion, 8 weeks old C57BL6/j mice were injected i.p. with LPS (Sigma L3755) at 150 μg/25 g body weight. DEXA (10 mg/kg) and TMs (40 mg/kg) were co-administrated with LPS via i.p. injection in a solution containing 20% Kolliphor and 5% DMSO in PBS. Blood samples were collected 90 minutes later by cardiac puncture, and two serial 10 min centrifugations, respectively at 3000 rpm at 4°C and at 6000 rpm, were performed to collect sera.

### Biotinylated RNA Pull Down assay

5-10 million RAW264.7 cells were seeded and treated with either DMSO or TMs (10 μM) for 6 hours, then lysed in polysome Extraction Buffer (PEB)(20 mM Tris-HCl (pH 7.5), 100 mM KCl, 5 mM MgCl_2_ and 0.5% NP-40 plus RNAse and protease inhibitors), and incubated for 1 hour at 4°C with 0.5 μM of positive (BiTNF-) or negative biotinylated probes^34^ (BiTNFneg, 5′-ACCACCCACCACCCACCCACCACCCA-3’) RNA in a TENT buffer (20 mM Tris-HCl (pH 8.0), 2 mM EDTA (pH 8.0), 500 mM NaCl 1% (v/v), Triton X-100 plus 100 units of RNAse inhibitors (Thermo Fisher Scientific) and protease inhibitors (Sigma Aldrich)^38^. Solutions were incubated for further 2 hours with 30 μL/samples of streptavidin magnetic beads (Life technologies, 11205D). 10% of the total lysates for each sample was kept and used as input material for subsequent precipitation assay. Specific protein enrichments in beads-precipitated samples were analyzed by immunoblotting and densitometric analysis obtained using the Image J 1.4 software (NIH). Samples were diluted in Laemmli Buffer (6X), denatured at 98°C for 5 minutes, and then separated by SDS–PAGE and blotted onto PVDF membranes (Immobilon-P, Millipore). Membranes were incubated for 1 hour at RT or overnight at 4°C with the following antibodies: mouse anti-HuR (Santa Cruz, sc-71290) and mouse anti-βACTIN antibody (3700, Cell Signaling). Secondary antibodies (Santa Cruz) were used for protein detection, using an ECL (Enhanced ChemiLuminescence) Select Western Blotting Detection Reagent (RPN2235; GE healthcare). Immunoblotting for β-actin was performed as control.

### Enzyme-linked immunosorbent (ELISAs) and Bioplex assays

ELISAs were carried out on mice sera, RAW264.7 and BMDMs supernatants with several dilutions according to each targeted cytokine. For Cxcl10 detection, RAW264.7 and BMDMs supernatants were diluted 1:5. Mice sera were diluted respectively 1:10 (CXCL10 detection) and 1:5 (TNF detection), according to manufacturer’s instructions (Mouse CXCL10/IP-10/CRG-2 DuoSet ELISA # DY466 and Mouse TNF-alpha DuoSet ELISA # DY410, R&D systems). Signals were detected by using TMB solution (Thermo Fisher Scientific) as a substrate. The reaction was then stopped with 2N H_2_SO_4_ and read with a Tecan microplate reader at 450 nm. Cytokine analysis was performed with the Bioplex technology (BioRad Laboratories), which combines a sandwich immunoassay with fluorescent bead-based technology, allowing individual and multiplex analysis of up to 100 analytes in a single microtiter well^39^. The assay for mouse Tnfα, Il1b, Il6, Il10 and Cxcl10 (Merck, Darmstadt, Germany) was carried out at Bioclarma srl, Torino, Italy. Briefly, serum samples were diluted 1:2 in assay buffer and analyzed in 96-well microplates, accordingly to the recommendations of manufacturers (Bio-Rad Laboratories Bio-Rad Laboratories, Hercules California, USA). The content of each well was then drawn up into the Bio-Plex 100 System array reader (Bio-Rad Laboratories Bio-Rad Laboratories, Hercules California, USA), which identifies and quantifies each specific reaction based on bead color and fluorescent signal intensity. The data were finally processed using Bio-Plex Manager software (version 6.1) using five-parametric curve fitting and converted in pg/mL.

### Immunofluorescence

2*10^4^ RAW264.7 cells/well were seeded in a 96-well cell carrier plate and after treatments (see Results) they were fixed with 3.7% paraformaldehyde (PFA) for 15 min at RT. Cells were permeabilized for 10 min with permeabilization buffer (0.2% Triton X-100 in PBS), and incubated with blocking solution (2% Bovine Serum Albumin in PBS) for 15 min. Primary antibody anti-HuR 1:250 in 3% BSA, anti-NF-kB 1:250 in 3% BSA and secondary fluorophore conjugated (Alexa 594 Red or Alexa 488) antibody (1:500) were diluted in PBS + BSA 0.6%. DAPI Blue (1.5 μg/mL) in PBS + BSA 0.6% was used to detect nuclei. Fluorescence images were acquired using ImageXpress Micro Confocal (Molecular Devices). In each well, images were acquired in 5 preselected fields of view with 20X Plan Apo objective (0,75NA) over three channels: Blue λEx= 377/54 nm - λEm= 432/36 nm, Green λEx= 475/28 nm - λEm= 536/40 nm, FarRed λEx= 635/22 nm - λEm= 692/40 nm. For optimal detection four z-stacks were acquired with a step size of 3µm in Spinning Disk Confocal mode (60 µm pinhole), the resulting maximum projection images were used for the Analysis. In brief, individual cell nucleus and cytoplasm were segmented using MetaXpress Custom Module Editor (MD), the ratio between nuclear and cytoplasmic signals was calculated for each cell and the mean value of the well was reported.

### Actinomycin D chase experiments

Transcription was blocked with Actinomycin D (ActD) administration at 2.5 uM respectively for 1 and 3 hours to measure mRNA stability. After experimental optimizations, ActD treatments were performed after 3 hours stimulation of LPS to guarantee inflammatory response activation. TM7nox (10 uM) was added simultaneously with ActD or with LPS (1 ug/mL) to evaluate its capability to modulate targets transcription or degradation/stability. Total RNA form each sample was extracted with TriZol and qRT-PCR were performed to quantify mRNA level as described above.

### Computational Details: Docking Calculations

The crystal structure of the RRM1 and RRM2 HuR domains complexed with m-RNA (PDB ID: 4ED5)^40^ was chosen as starting receptor conformation for docking simulations. The choice of the “closed” form, due do the presence of mRNA, was based on the capability of our TMs to bind HuR in the mRNA binding region. Both HuR and the ligands were prepared with Maestro^41^ Version 12.7.156, the interface for Schrödinger’s molecular modeling platform. In particular, the protein was prepared with Protein Preparation Wizard^42,43^, included in Maestro. Hydrogens were added to the protein, and missing side chains were added by using the Prime^44–46^ module of Maestro. Crystallographic water molecules and the native ligand m-RNA were deleted. The N-terminal and C-terminal residues were capped with acetyl (ACE) and N-methyl amide (NME) groups, respectively. To properly describe the protonation state of the protein residues and also correctly describe the hydrogen bonding networks at pH=7, protonation states were assigned by evaluating their pKa with the Propka^47^ program included in Maestro. Finally, a relaxation procedure was performed by running a restrained minimization only on initially added hydrogen atoms, according to the OPLS2005^48^ force field. TM ligands were generated and then prepared through the LigPrep^49^ module of Maestro, employing the OPLS2005 force field. The Epik^50–52^ module of Maestro was used to evaluate the pKa of each ligand at pH=7, to properly describe its protonation state. Each obtained TM ligand was further optimized at molecular mechanics’ level through the MacroModel^53^ program included in the Schrödinger suite of programs. Docking simulations were performed with Autodock4.2^54^. The grid and setting to run docking calculations were prepared by using AutodockTools, the graphical interface of Autodock. The grid for docking calculations was computed by using 108×108×68 points, spaced by 0.375 Å. These parameters correspond to a grid of ∼40x∼40x∼25 Å in a three-dimensional Cartesian coordinate system. In so doing, the whole region between the two HuR domains in the XY-plane was properly included. A hundred independent runs of the Lamarckian genetic algorithm local search method per docking calculation were performed, and a threshold of maximum 25 million energy evaluations per run was applied. Docking conformations were clustered on the basis of their RMSD (tolerance = 2 Å).

### Molecular Dynamics Simulations

The force field ff14SB^55^ was used to model HuR. Regarding **TM7nox**, the Generalized Amber Force Field^56^ was employed. RESP^57^ charges were obtained by using the antechamber accessory module of AmberTools. More in detail, the ESP^58^ charges employed to calculate RESP ones were evaluated at *ab initio* theory level with the Gaussian^59^ software. The TM was optimized by using the density functional theory method, that has accurately simulated molecular structural and spectroscopic properties^60–65^. In particular, the B3LYP^66^ /6-31G* theory level was employed, and then ESP charges were calculated on the optimized minimum energy structure at HF^67^/6-31G* theory level. By using the leap program available in AmberTools, a solvent box of 12 Å between any protein atom and the edge of the box was added, where water molecules were described through the TIP3P^68^ force field. A box of about ∼70x∼65x∼85 Å was obtained, while neutrality was ensured by adding six Cl^-^ ions, modeled with Joung and Cheatham^69^ parameters. Finally, coordinates and topology files for the whole system were obtained. Energy minimizations and MD simulations were performed with Gromacs^70,71^ software. For all simulations (both equilibration and production runs), the Verlet cut-off scheme was used for non-bond interactions neighbor search. The Fast smooth Particle-Mesh Ewald^72^ (SPME) method was employed for long-range electrostatic interactions; the cut-off was set to 1.2 nm for long-range Van der Waals interactions, with the Lennard-Jones potential gradually switching to zero between 1 and 1.2 nm. For all MD simulations, the leap-frog^73^ algorithm for integrating Newton’s equations of motion was used and a time step of 2 fs was chosen. The equilibration procedure started with two energy minimization steps performed with the steepest descendent algorithm. The first one was 20000 steps long, the ligand and the protein heavy atoms were kept fixed by imposing a harmonic constraint of 500 kcal mol^-1^ Å^-2^, so that only the solvent was allowed to relax. In a second, 10000 steps long equilibration run the entire system was not constrained. Then, the system was gradually heated by increasing the temperature by 50 K in each step, with subsequent MD runs in the canonical ensemble (NVT), until reaching a final temperature of 300 K. All these steps were 200 ps long, and harmonic restraints were gradually decreased in each step. Beside the first one, constraints of 30 Kcal mol^-1^ Å^-2^ were applied on all the heavy atoms of the protein and the ligand; for all the other NVT simulations constraints of 25, 18, 12, 6.5 Kcal mol^-1^ Å^-2^ were applied only on the protein backbone atoms (Ca, N, C, O) and the ligand heavy atoms. During the last NVT step a constraint of 1.5 Kcal mol^-1^ Å^-2^ was applied only on the ligand. The weak-coupling Berendsen^74^ scheme was used for temperature coupling. A final NPT run of 1 ns was performed without constraints, to adjust the box volume, and the Berendsen algorithm was used for pressure coupling. MD production runs of 2 µs with a time step of 2 fs were performed for each of the chosen poses. For production runs, temperature and pressure controls were carried-oud with the velocity-rescale^75^ and Parrinello-Rhaman^76^ scheme, respectively. The LINCS^77^ algorithm was employed to constrain bonds. Trajectory visualization and analyses were performed with the VMD^78^ software, while the figures were obtained using the Pymol^79^ molecular visualization system.

### RNA Immunoprecipitation and NGS-sequencing

RAW 264.7 cells (30 million) were used in RIP experiments, followed by qRT-PCR or NGS sequencing. In general, RIPs were performed without cross-linking steps^80^, using 1-15 μg/mL of anti-HuR antibody (Santa Cruz, 71290) and the same amount of mouse normal IgG isotype (negative control, Santa Cruz 2025). Cells were harvested after treatments for 6 hours with LPS (1 μg/mL), TMs (10 μM) or DMSO, then lysed with 20 mM Tris–HCl at pH 7.5, 100 mM KCl, 5 mM MgCl_2_, and 0.5% NP-40 supplemented with 1% protease inhibitor cocktail (Sigma Aldrich) and 100 units of RiboLOCK RNase inhibitor (Thermo Fisher Scientific) for 10 min on ice, and centrifuged at 15.000 × g for 10 min at 4°C. Lysates were incubated with Pierce A/G beads (Thermo Scientific Pierce 88847-88848) for 1 hour at 4°C for pre-clearing steps, and in parallel 80% A and 20% G beads for each sample were incubated either with HuR or IgG antibodies for Ab-coating step for 1 hour at RT. Then, lysates were incubated with antibodies and beads for further 4 hours at 4°C. Finally, samples were washed (6 times, 5 minutes each wash) with NT2 buffer. The TRIzol reagent was added directly to the beads for HuR-bound RNA isolation, and they were processed for qRT-PCR analysis or library preparation. 1-5% of the total lysate for each sample was stored as input. For validation experiments, quantitative PCRs were performed after cDNA synthesis (Thermo Fisher Scientific, K1612) using Universal SYBR Master Mix (KAPA Biosystems, KR0389) on CFX-96/384 thermal cyclers (BIO-RAD). Fold enrichment for *Cxcl10, Cd40, Fas, Nos2, Il1bβ* was calculated as previously described^35^. In detail, we applied the equation 2e-ΔCt, in which ΔCt is expressed as the ratio between target mRNA IP HuR on target mRNA IgG. For each condition, the ΔCt value for HuR and IgG IP samples were calculated in triplicate.

### cDNA Library preparation

Quantity and quality of RNA samples (RNA-HuR IP; IgG IP and inputs) were measured by using a Qubit− RNA high sensitivity (HS), broad range (BR) (Thermo Fisher Scientific). RNAs from INPUT and RIP were quality-controlled using Bioanalyzer (Agilent technologies) and 7 ng of the extracted RNA, having RIN higher than 9, were used for fragmentation at 94 °C for 4 min. RNA libraries were generated using the SMART-Seq Stranded Kit (Takara). The kit incorporates SMART® cDNA synthesis technology^81^ and generates Illumina-compatible libraries via PCR amplification, avoiding the need for adapter ligation and preserving the strand orientation of the original RNA. The ribosomal cDNA was depleted by a ZapR-mediated process, in which the library fragments originating from rRNA and mitochondrial rRNA are cleaved by ZapR in the presence of mammalian-specific R-Probes. Library fragments from non-rRNA molecules were enriched via a second round of PCR amplification using Illumina-specific primers. Quantity and quality of each individual library were defined using a Qubit Fluorometer (Thermo Scientific) LabChip GX (Perkin Elmer). After libraries’ equimolar pooling, the final pool was quantified by qPCR (KAPA and BIORAD). The adaptor-tagged pool of libraries was loaded on 1 NovaSeq6000 SP flowcell (PE100 chemistry) for cluster generation and deep sequencing, producing an average of 40M reads per sample (Illumina, San Diego, CA).

### Bioinformatics Analysis

RNA sequencing and RIP sequencing were performed in quadruplicate (GEO number GSE198605). Raw sequence files’ quality was checked via FastQC (v 0.11.9, http://www.bioinformatics.babraham.ac.uk/projects/fastqc/). Gene quantification was conducted with STAR (version v. 2.7.7a) starting from Ensembl GRCm39 genome version. The generated genes counts were analyzed using DESeq2 package. The normalized count matrix (obtained from variance stabilizing transformation (VST) method as implemented in DESeq2 package) was used to explore high-dimensional data property with Principal Component Analysis (PCA) coupled with a dimensionality reduction algorithm used in the DESeq2 package. Differentially expressed genes (DEGs) were selected with a *p-adjusted* cut off of 0.05 and a log2 Fold Change value greater than 1 (up-regulated DEGs) or lower than -1 (down-regulated DEGs). *P-value* was adjusted for multiple testing using the Benjamini–Hochberg (BH) correction with a false discovery rate (FDR) ≤ 0.05. DEGs were then analyzed with a hierarchical clustering method, using correlation distance. Visualization of Z-score normalized values and clustering was obtained via pheatmap package, while visualization of DEGs in volcano plots were acquired using the EnhancedVolcano package. Venn diagrams were created using vennPlot (systemPipeR package). Barplots of genes of interest were created using ggbarplot, annotate_figure, ggarrange (ggpubr package), add_pval (ggpval). Functional annotation was performed for all the comparisons and for gene list of interest. We used both clusterProfiler and Reactome Bioconductor packages, whose results were visualized with ggplot2 and ggpubr packages. The full enriched annotation table results are provided in the Supplementary files. A treeplot and two different networks of the most important pathways resulting from the functional annotation analyses were generated using the Bioconductor package enrichplot.

The correlation analyses among RNAseq and RIPseq shared genes was performed using cor.test function and, for better visualization, we subtracted the two correlation data and plotted the result in a 3D representation, generated with a Delaunay triangulation and a Dirichlet tessellation. In the analysis of the properties of the 3’-UTRs (the lengths and U/AU-rich regions (ARE)) between the differential genes in the TM7nox and LPS co-treatment versus LPS and DMSO RIP-seq, we also included the not differential genes as a control. The length of the 3’-UTR were calculated from the 3’-UTR sequences downloaded from Ensembl (accessed in August 2022). The ARE was downloaded from the Database for AU-rich elements and direct evidence for Interaction and lifetime regulation (Aresite2, http://nibiru.tbi.univie.ac.at/AREsite2 accessed in August 2022). These data were visualized via boxplots and statistical support was provided by a wilcoxon test. Ten HuR binding siteswere obtained from EuRBPDB, a database for eukaryotic RNA binding proteins containing 315000 RBPs from 160 species^82^.

### Total RNA extraction and qRT-PCR

Total RNA was extracted either with a RNA extraction kit (Zymo Research), according to the manufacturer instructions or with Trizol, chloroform precipitation followed by RNAse free-DNAse I treatments, 15 min at 37°C. cDNA Synthesis (Thermo Scientific, K1612) was performed from 1 μg of RNA, and qRT-PCRs were performed using Universal SYBR Master Mix (KAPA Biosystems, KR0389) on CFX-96/384 thermal cyclers (BIO-RAD)^30,31,35^. Normalized expression levels for each selected gene were calculated as 2e-ΔΔCt, where the Ct value of either control or treatment conditions was subtracted from the Ct value of the housekeeping genes (RPLP0) to yield the ΔCt value. Then, the ΔCt value for treatment and control were computed in duplicate and averaged to give one ΔΔCt value per sample.

### Statistical analysis

Statistical analysis experiments were performed in a number of biological replicates indicated in all experiments reported in the Results section. t-tests were used to calculate final p-values, without assuming variances to be equal (Welch’s t-test). P-value <0.05, < 0.01, <0.001 and <0.0001 were indicated with *, **, ***, **** symbols, respectively.

## Results

### TMs show a redox keto-enolic tautomerism

In earlier efforts, we identified naturally occurring dihydrotanshinone I (DHTS, **1**)^31,35^ and a DHTS-inspired family of synthetic tanshinone mimics (TMs), such as unsubstituted TM **2** / **TM6a** (Figure 1A)^30^. We selected our most potent earlier TM **3** / **TM6n**, (Figure 1A), and we aimed to introduce two orto substituents on the 3-phenyl ring of the 3-aryl-1,3-aza tanshinone system (**4ox** / **TM7nox**, Figure 1A), to possibly increase solubility as planarity disruption is known to lead to higher solubility of orto-substituted compounds^83^. Furthermore, we explored alternative routes to higher bioavailability by reducing the orto quinone to a more hydrophilic orto diphenolic compound (**4red, TM7nred**), and we acetylated the latter to yield a putative esterase-sensitive prodrug **5 / TM8n** which could escape any quinone-driven metabolic instability before reaching its molecular target.

**Figure 1.**
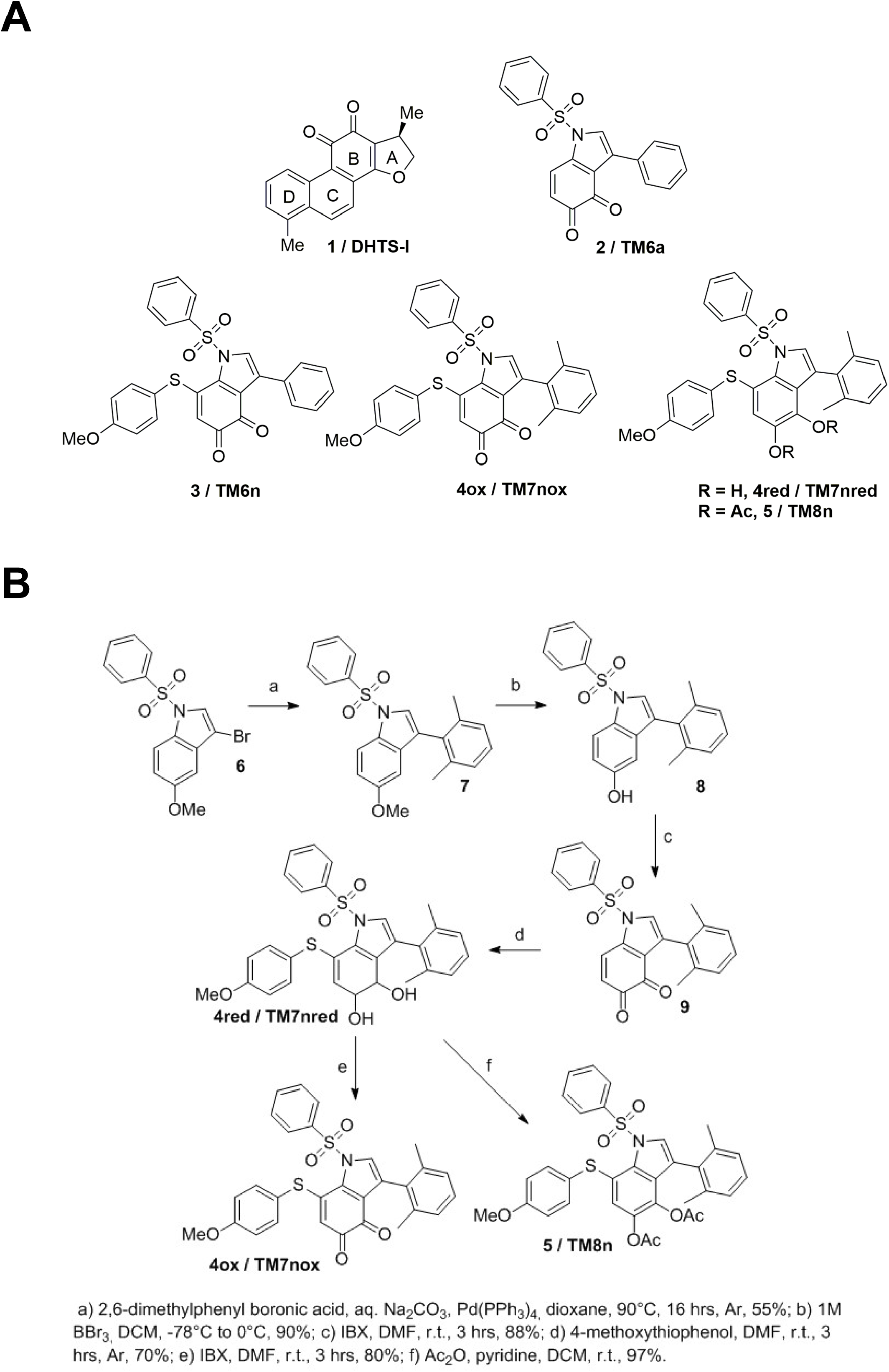
Chemical structure and synthesis of TMs. A) Chemical structure of standard DHTS-I 1 and unsubstituted **TM6a 2** (top); of early lead **3** / **TM6n**, of dimethyl quinone **4ox** / **TM7nox**, dimethyl diphenol **4red** / **TM7nred** and dimethyl diacetate **5** / **TM8n**. B) Synthesis of dimethyl quinone **4ox** / **TM7nox**, dimethyl diphenol **4red** / TM7nred and dimethyl diacetate **5** / **TM8n**.

Compounds **TM6n** (**3**), **TM7nox** (**4ox** - quinone), **TM7nred** (**4red -** diphenol) and **5 / TM8n** were synthesized and profiled in biological and physic-chemical assays. Working on the published synthesis of **TM6n**^30^, we improved it to yield bis-orto substituted TM **4** / **TM7nox, TM7nred** and **5** / **TM8n** (Figure 1B) on a multi-gram scale. A standard Suzuki coupling protocol using 2,6-dimethylphenyl boronic acid led to desired 3-aryl 5-methoxy intermediate **7** (Figure 1B) in poor yields (∼6%), likely due to the additional steric hindrance around the reaction site. Conversely, by switching to dioxane as a solvent and raising the reaction temperature (Figure 1B, step a) we obtained target 3-aryl 5-methoxy intermediate **7** in moderate yields. Standard demethylation and oxidation of phenoxy intermediate **8** (Figure 1B, steps b and c respectively) led to 3-(2,6-dimethyl)-phenyl TM **9** in good yields. The reduced, diphenolic 7-functionalized target **4red** / **TM7nred** was then obtained in good yields following a standard 1,4-Michael addition protocol on intermediate **9** (step d^30^). A portion of diphenolic **4red** was then oxidized to orto-quinone **4ox** / **TM7nox** using a standard IBX protocol (step e), while another portion was easily converted into the acetylated pro-drug derivative **5** / **TM8n** by almost quantitative acetylation (step f, Figure 1B).

We then investigated the **TM7nox/TM7nred** interconversion equilibrium – in particular, the ability of diphenolic **TM7nred** to spontaneously convert to quinonic **TM7nox** in various media. ^1^H-NMR studies were performed, observing that diphenolic **TM7nred** reached an equilibrium at RT with quinone **TM7nox** when dissolved in organic solvent (acetone-d_6_), reaching ∼35% conversion into the oxidized form in 48 hours (Supplementary Figure S1A). The conversion rate of **TM7nred** into **TM7nox** increased when D_2_O (10% v/v) was added to the solvent, reaching ∼46% after 24 hours (Supplementary Figure S1B). Conversely, the oxidized form **TM7nox** dissolved in acetone-d_6_ showed higher stability and limited, ∼12 % conversion into diphenol **TM7nred** after 45 days/1.5 months. Thus, we can safely assume that the biologically active/HuR-binding orto-quinone TM form should be the major species in biological environments, in agreement with modeling data.

We then considered an alternative pro-drug approach^84^ to prevent metabolic transformations or off target interactions due to such redox equilibration. In particular, by masking the orto-quinone of Michael adduct **TM7nox** as a diacetate (**5** / **TM8n**, Figure 1A) we aimed at obtaining a putative prodrug incapable of binding to HuR that can be converted *in situ* to reduced **TM7nred** by esterases, after reaching its cellular or *in viv*o targets. This reduced TM – as determined by preliminary redox equilibration studies reported above – should eventually be converted into biologically active quinone **TM7nox** after O_2_-promoted equilibration in cellular media.

Early lead **TM6n**, the redox couple **TM7nox** and **TM7nred**, and diacetate **TM8n** were tested to determine their solubility in suitable aqueous media for *in vitro* and *in vivo* testing. We initially planned kinetic solubility measurements (LC-MS) in PBS buffer with increasing quantities of biocompatible solubilizer-excipient Kolliphor EL; unfortunately, the latter resulted to be unsuitable, as its UV spectrum covers the peaks of our TMs and prevents their quantitation. Thus, we replaced Kolliphor EL with DMSO, and we surprisingly characterized early lead **TM6n** as the most soluble TM (from 0.8 μM in PBS-5% DMSO to 4.2 μM with 20% DMSO), followed by **TM7nred** (from 1.5 μM in PBS-10% DMSO to 2.9 μM with 20% DMSO), **TM7nox** (from 0.41μM in PBS-10% DMSO to 1.8 μM with 20% DMSO) and **TM8n** (0.2 μM in PBS-20% DMSO). A graphic summary of kinetic solubility up to 20% DMSO for **TM6n, TM7nred, TM7nox** and **TM8n** is provided (Supplementary Figure S2). Although such results were discouraging, we attempted to solubilize the TMs in the medium for *in vivo* testing (PBS buffer, DMSO 5%, Kolliphor EL 20%), and we observed a significantly different behavior. Namely, dimethylated **TM7nred, TM7nox** and **TM8n** could be completely dissolved, while **TM6n** could not as it precipitated in the above-mentioned buffer. Therefore, we decided to further characterize our TMs, while prioritizing **TM7n**-related (**TM7nred, TM7nox** and **TM8n**) derivatives for *in vivo* administration.

### The ortoquinone TM7nox is the active form of TM7 series and keeps HuR in a closed conformation

Our earlier results show that DHTS (**1**) and our TMs molecules, such as **TM6a**, bind to the β-platforms of the RRM1 and RRM2 domains, stabilizing a HuR closed conformation that hampers the accommodation of the binding mRNA^30^. However, **TM7n** derivatives (**TM7nred, TM7nox, TM8n**) are more sterically hindered. Moreover, the planarity disruption induced by two orto-methyl substituents on the 3-phenyl ring could partially change its stacking surface with respect to the previous synthesized TMs, so that a different accommodation within HuR must be considered. Thus, we investigated the molecular binding mode of **TM7nox**, the active and prevalent molecular species in solution. Molecular docking experiments were carried out by using AutoDock4.2 (see Materials and Methods), allowing **TM7nox** to explore the entire interdomain space. Given the large region explored and the smaller size of **TM7nox** compared to mRNA, AutoDock did not find a unique binding mode. Thus, we focused on the lowest energy pose and on those representing the most populated clusters, referred from now on as PI, PII and PIII, respectively. All three poses occupy the RRM1-RRM2 interdomain region and differ by a diverse closeness to the hinge region (Figure 2A). Particularly, as to PI, **TM7nox** is placed closer to the hinge loop, resembling the binding mode found for **TM6a**^30^. As to PII and PIII, **TM7nox** occupies the RRM1-RRM2 inter-domain region in a more central area (less close to the hinge loop). To assess the relative stability of each pose, 2 µs long MD simulations were then performed. The time evolution of **TM7nox** during the trajectories is graphically represented in Figure 2B, where the **TM7nox** center of mass is shown as a sphere colored according to the simulation time (from red-initial poses to blue-final poses), while the plots of the RMSD as a function of time with respect to the first step are shown in Supplementary Figure S3. The molecular dynamics simulation of PI shows that **TM7nox** remains in the hinge region, although it changes its orientation during time because of the flexibility of the loop, that makes difficult to retain stable interactions with neighbor residues. Indeed, the position of the center of mass during the simulation suggests that **TM7nox** explores different possible arrangements in the hinge loop (Figure 2B). Despite this, the ligand stays in this binding region for the entire trajectory, as clearly shown also by the RMSD plot (see Supplementary Figure S3).

**Figure 2.**
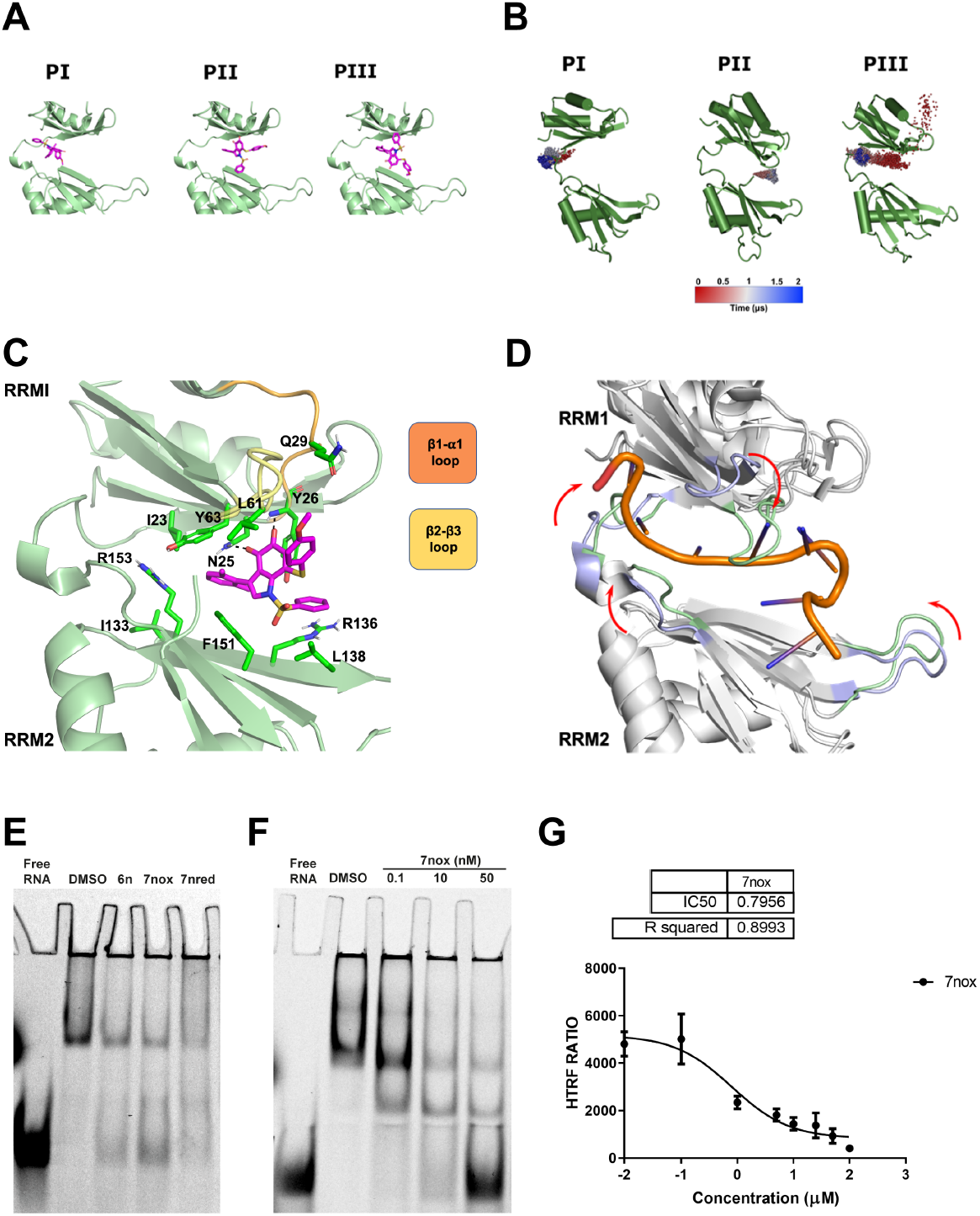
TM7nox binds to HuR and disrupts HuR RNA binding ability *in vitro*. A) Ligand binding poses found by Autodock4.2 and submitted to MD simulations. The ligand is schematically shown in purple sticks and HuR in green cartoon representation. B) Representation of the **TM7nox** exploration of the HuR cavity for each simulated pose. HuR is represented as green cartoon while the ligand center of mass evolution during the trajectory is shown as colored sphere. C) Theoretical **TM7nox** binding mode-PII, as suggested by our MD simulation. The ligand is shown in purple sticks and the protein as green cartoon. Main residues involved in interactions with the ligand are shown as green sticks. Nonpolar hydrogens are hidden for clarity. D) Superposition of the HuR-RNA complex crystal structure with the final state of the dynamized PII. The secondary structures are depicted in grey cartoons while the loops in light blue for the initial state and light green for the last frame of the MD simulation. E) Representative REMSA showing HuR-RNA binding impairment induced by TMs. rM1M2_HuR [3.7 nM] was incubated for 30 min with either 1 nM of 5′-DY681-labeled RNA probe alone, or together with DMSO used as control and **TM6n, TM7nox** and **TM7nred** at 1 μM doses. F) Representative REMSA showing **TM7nox** dose-response inhibition of the binding between 100 nM of rM1M2_HuR and 1 nM 5′-DY681-labeled RNA probe. G) Concentration response analysis of **TM7nox** tested in HuR:RNA probe interaction assay. In a dose dependent manner **TM7nox** [3,125 μM; 6,25 μM; 25 μM; 50 μM; 100 μM; 150 μM; 200 μM] interferes with the binding between His-tagged recombinant M1M2 HuR protein [20 nM] and 5’Bi-TNF ARE probe [50 nM]. The calculated IC_50_ is 0.7956 μM and data have been normalized on control (DMSO). Data fit nonlinear regression fitting curves according to a 1-site binding model in GraphPad Prism. Plotted bars are mean ± SD of three independent experiments

Similarly, in the PIII case, **TM7nox** once more accommodates in the hinge loop region, trying to establish stable contacts with the loop residues, after an important ligand reorientation (after about 500 ns). Conversely, the PII pose seems to achieve clear and stable receptor-interactions (Figures 2B, 2C). In fact, the orto quinone oxygens are involved in two tight hydrogen bonds with the N25 side chain amide and with the Y26 NH backbone (see Supplementary Figure S4). Moreover, a “cage” of aromatic residues (Y26, Y63 and F151) locks the central indole ring position during the simulation (see Figures 2C and Supplementary Figure S5). As for the dimethyl-phenyl moiety, it is buried in the interdomain cavity and is oriented toward the hinge loop, establishing an amide-π interaction with N25 and other hydrophobic contacts with Y63, I23, I133, F151 and R153 residues. The phenylsulfonyl and 4-methoxyphenylthio substituents rearrange themselves so that the former is kept in place for the whole simulation time by a strong cation-π interaction with R136 and other hydrophobic contacts within the RRM2 β-barrel (F151 and L138); the latter plunges between the β2-β3 and the β1-α1 loops of the RMM1 domain, contacting the L61 and Q29 residues. Noteworthy, looking at the superposition of the HuR-RNA complex crystal structure on the final state of the dynamized PII pose (Figure 2D), the secondary structures are basically conserved during the simulation, while the loops surrounding the RNA binding cavity fold around the ligand, in order to dramatically reduce the available buried surface area between the two RRM domains. In particular, the β2-β3 loop flaps in the region where the A7-RNA base is located (Figure 2D). Thus, in the PII final pose **TM7nox** stabilizes HuR in a conformational state that hampers the accommodation of a binding RNA sequence. Our results suggest that for **TM7nox**, although binding modes close to the HuR hinge loop are plausible and were found by both docking and 2 µs MDs, the center of the cavity made up by the beta-sheets of the two RRM domains is the only receptor region to establish stable contacts with the ligand moieties. In fact, the most significant difference between the PI-PIII binding modes and PII is, indeed, the lack in the former poses of two permanent H-bond interactions established by the quinone group. Finally, in line with experimental data (see Supplementary Figure S6A), molecular docking demonstrated that the binding of **TM8n** (prodrug-inactive derivative of **TM7nox**) to HuR is strongly unfavored. In fact, all its binding poses are higher in energy in comparison with **TM7nox**, and there is a failure in finding a convergence among the obtained poses (most clusters are populated by a single pose), suggesting that it is hard to find a reliable accommodation for **TM8n**. In summary, due to chemical differences between **TM7nox** and **TM6a**, they bind HuR in a slightly different mode. Nonetheless, molecular modeling strongly suggests that they share the same mechanism of action, stabilizing an HuR closed conformation unable to accommodate the mRNA.

### Orto-dimethyl TMs interfere with the HuR-RNA complex

The first two HuR domains were recombinantly (rHuR) produced in *E*.*Coli BL21* and incubated with a 26 nt long, ARE containing RNA probe linked in 5′ to the DY681 fluorophore to form the protein-RNA complex. REMSAs (RNA Electro Mobility Shift Assay) revealed that **TM6n, TM7nox, TM7nred** and **TM8n** interfere with the binding of the protein with the RNA probe while non-denaturing gel electrophoresis does not suggest the inhibition of protein dimerization (Figure 2E, 2F and Supplementary Figure S6A-B). Such results are coherent with computational estimations and analytical studies, due to the expected compatibility of orto-dimethyl substitutions on the **TM6n** scaffold with HuR binding; and with the redox quinone-diphenol equilibrium, providing also reduced **TM7nred** with the ability to disrupt the HuR protein-RNA complex via its conversion to **TM7nox** in biological media and the expected negative effect of the diacetate functionalization on HuR binding. To measure the disrupting ability of HuR:mRNA complex by **TM7nox**, we set up a HTRF (Homogenous Time-Resolved Fluorescence) assay. We used biotinylated 26 nt ssRNA probes containing the AU rich elements of the TNFα Bi-TNF to determine the hook point of the assay^34^ (Supplementary Figure S6C). We then titrated the protein-RNA complex with increasing concentrations of **TM7nox** and obtained an IC_50_ of 0.79 µM (Figure 2G). Then, the activity of **TM6n** and **TM7** derivatives in counteracting HuR were investigated in macrophages during LPS stimulation, a model in which HuR is known to regulate the cell response^85–87^.

### TM7nox reduces inflammatory and chemotaxis response induced by LPS in murine macrophage RAW 264.7 cell line

We evaluated TMs’ toxicity in RAW 264.7 murine macrophage cell line, and in Bone Marrow Derived Macrophages (BMDMs) to better predict TMs effects *in vivo*. BMDMs were harvested from 6 to 12 weeks old C57BL6/j wild type mice and, after one-week *in vitro* differentiation of the monocytes into macrophages with L929 supernatant^36^, they were treated with **TM6n, TM7nox, TM7nred** and **TM8n** for 24 hours. TMs seem to show higher toxicity in primary cultures than in RAW 264.7 cells, considering that their IC_50_ is lower than 2 µM (Supplementary Figure S7A, B). To investigate the ability of TMs to modulate the macrophage response to inflammatory stimuli, we challenged RAW264.7 cells with LPS (1 µg/mL) for 6 hours. We chose **TM7nox** as a reference compound to evaluate its ability to counteract LPS induced response, as the orto quinone form is the one interacting with HuR. We measured the abundance of transcripts after LPS treatment, and the ability of **TM7nox** to disrupt the interaction of HuR with its target mRNAs by employing a transcriptome-wide approach of RNA preparations from untreated cells, LPS-treated and LPS+**TM7nox**-treated cells. Principal component analysis (PCA) shows that principal component 1 (PC1) and principal component 2 (PC2) explained most of data variability, with 83% of variance associated to PC1 and 7% of variance associated with PC2. The effect of LPS treatment on transcriptome changes separating LPS-untreated from LPS-treated samples can be appreciated along the PC1 axis. LPS-treated groups segregated along PC2, which instead identified the effect of **TM7nox** on LPS treated cells (Figure 3A). Therefore, LPS is the major modulator of gene expression changes but, within this context, **TM7nox** modulates the LPS cell response to a significant extent. Indeed, LPS triggered a strong response by inducing 2829 DEGs with respect to the vehicle (DMSO; 1354 logFC < -1, 1475 logFC > 1, Supplementary Table 1). Co-treatment of LPS with **TM7nox** still induced a strong response, as evaluated by the number of DEGs (3273, 1654 downregulated, 1619 upregulated, Supplementary Table 1). The number of genes commonly regulated by LPS, and by LPS-**TM7nox** co-treatment accounted for (1045 + 45) upregulated + (967 + 46) downregulated genes (Venn diagram, Figure 3B, Supplementary Figure S8A). This ensemble of genes is modulated coherently in both treatments, represents the strong majority of DEGs, and can be considered as an LPS-induced response that is not affected by **TM7nox**. The strong effect of LPS can be also observed in the heat-map of DEGs, in which unsupervised clustering clearly separated LPS-treated samples from control samples. However, **TM7nox** samples organized as a subcluster within the larger LPS branch (Figure 3C). Functional annotation identified several Gene Ontology (GO) terms related to the inflammatory process as the response to LPS, interferon *γ* and *β*, NF-kB activation and chemotaxis among upregulated genes; whereas among the downregulated genes we found several GO terms related to DNA replication and cell cycle (Supplementary Table 2). Among the top upregulated DEGs we found *Cxcl10, Fas, Il1b, Cd40, Nos2, Cxcl2, Il6* and *Il10* (Figure 3D). These pathways and genes are widely recognized as canonically activated by LPS in RAW264.7 cells, validating our experiment^86,88^. Three subgroups of regulated genes could also be identified (Venn diagram Figure 3B), i.e. genes regulated by LPS/**TM7nox** vs DMSO, by LPS vs DMSO and by LPS/**TM7nox** vs LPS (Supplementary Table 1). We focused our attention on the last subgroup, i.e. emerging categories from upregulated DEGs modulated by **TM7nox** in presence of LPS (DEGs: 170). They provide the enrichment of cation transmembrane transporter activity genes (Supplementary Table 3), while among the downregulated genes (DEGs: 249) we observed a strong enrichment of categories related to the inflammatory response, cytokines (*Il1b, Cxcl10, Il10, Il19, Il33*), immune cell chemotaxis (*Ccl12, Ccl22, Ccl17, Ccl6*) and innate immune response (Supplementary Table 3, Supplementary Figure S8B, C). The top five ranking GO pathways for the 249 down-regulated genes of interest were further visualized with a tree-plot (Figure 3E) and network of the present DEGs (Figure 3F). These results indicate that LPS-induced cellular response is not abolished by **TM7nox**, but rather modulated and mitigated. Specifically, **TM7nox** modulates the inflammatory response, downregulating the expression of important cytokines (*Il1b, Cxcl10*) and mainly influencing the expression of genes involved in cell chemotaxis. The modulation of LPS-induced response is further witnessed by the presence of an exclusive LPS response that is absent during **TM7nox** co-treatment.

**Table 1:** List of 20 DEGs resulting from analysis of TM7nox/LPS co-treatment and DMSO/untreated.

| **Gene name** | **Description** |
| --- | --- |
| Il10 | Interleukin 10 |
| Cd40 | CD40 antigen |
| Tnfaip3 | tumor necrosis factor, alpha-induced protein 3 |
| Nos2 | nitric oxide synthase 2, inducible |
| Acod1 | aconitate decarboxylase 1 |
| Fas | Fas (TNF receptor superfamily member 6) |
| Il1b | interleukin 1 beta |
| Il1a | interleukin 1 alpha |
| Tnfrsf8 | tumor necrosis factor receptor superfamily, member 8 |
| Cxcl9 | chemokine (C-X-C motif) ligand 9 |
| Ccl22 | chemokine (C-C motif) ligand 22 |
| Ccl17 | chemokine (C-C motif) ligand 17 |
| Cxcl10 | chemokine (C-X-C motif) ligand 10 |
| Ccl12 | chemokine (C-C motif) ligand 12 |
| Ccl7 | chemokine (C-C motif) ligand 7 |
| Ccl2 | chemokine (C-C motif) ligand 2 |
| Thbs1 | thrombospondin 1 |
| Zc3h12a | zinc finger CCCH type containing 12A |
| Il27 | interleukin 27 |
| Cxcl11 | chemokine (C-X-C motif) ligand 11 |

**Figure 3.**
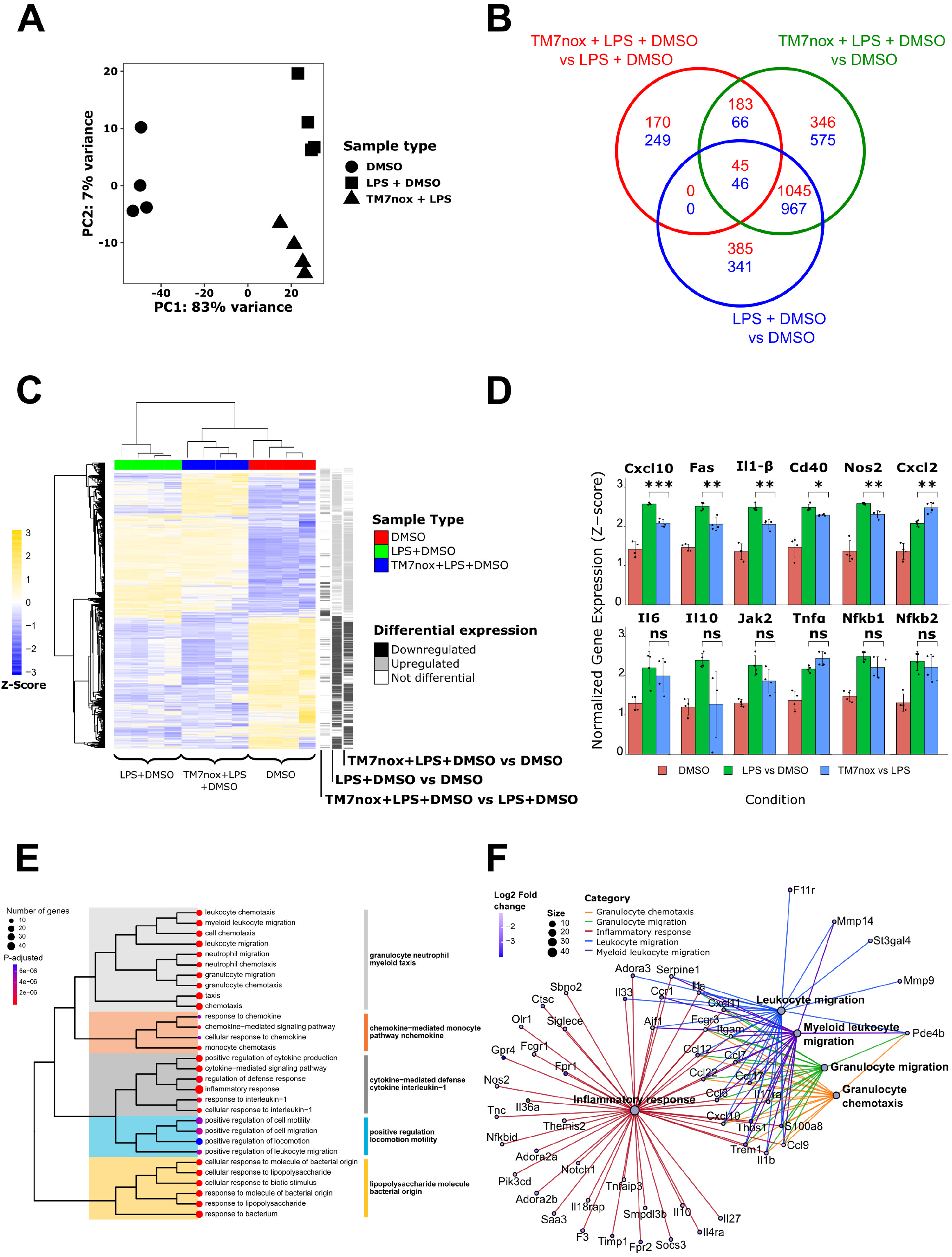
RNA-seq analyses reveals that TM7nox modulates LPS induced response. A) Principal Component Analysis (PCA) of the 12 samples. PC1 shows 83% variance and PC2 7%. Each dot represents a DMSO sample, each triangle is a sample treated with DMSO + LPS, while each square is a DMSO + LPS + **TM7nox** treated sample. Every condition groups together with same type samples and it can be observed that the effect of **TM7nox** separates DMSO+LPS from DMSO+LPS+**TM7nox** conditions. B) Venn of differentially expressed genes (DEGs), the numbers in each circle represent the number of DEGs between the different comparisons while the ones overlapping are for mutual DEGs (DMSO+LPS+**TM7nox** vs DMSO+LPS in red, DMSO+LPS+**TM7nox** vs DMSO in green, DMSO+LPS vs DMSO in blue). C) Heatmap of Z-score of differential genes across different samples, each grouping together with its own sample type (red: DMSO; green: DMSO + LPS; blue: DMSO + LPS + **TM7nox**). The track on the right shows differential gene expression between different comparisons (black: down-regulated DEGs; grey: up-regulated DEGs; white: no changes in DEGs). “Average” was used as clustering method and “correlation” for clustering distance of both rows and columns. D) Barplot of Z-score of key genes across different samples (red: DMSO; green: DMSO + LPS; blue: DMSO + LPS + **TM7nox**). Significance between DMSO+LPS and DMSO+LPS+**TM7nox** samples is shown by an asterisk for each gene, along with standard error represented by error bars. E) Tree-plot of enriched terms deriving from the 249 down-regulated genes. Each dot represents an enriched term which is also colored according to p-adjusted values, spanning from red to blue. Terms’ dimensions are relative to the number of genes found to be enriched in that category. The subclusters and their names, visible on the right, are highlighted with a specific color. F) Network visualization of the top five ranking GO pathways for the 249 down-regulated genes. Each pathway is represented by a gray dot and highlighted with a peculiar color, each gene is connected to the pathways it belongs to. Color of the genes are illustrated from blue to white according to their log2 Fold Change values, while size of the pathways dots depends on the number of genes enriched for the pathway itself.

### TM7nox disrupts HuR interaction with selected AU rich containing transcripts

We performed a HuR RIP-seq (ribonucleoprotein immunoprecipitation followed by sequencing) experiment in RAW264.7 cells to appreciate the modulation of gene expression response induced by **TM7nox** on LPS treatment, and its dependence on its biochemical activity as a disruptor of HuR/RNA binding. The enrichment fold change was calculated by comparing HuR-bound transcripts in each condition. We found a strong positive association of transcripts bound by HuR after LPS treatment (3887) in comparison to DMSO associated transcripts (Supplementary Table 1)^31^. PCA indicated that cellular response to LPS strongly modifies the HuR-bound transcriptome, segregating LPS-treated versus untreated samples and justifying the observed 86% variance. PC2 described the effect of **TM7nox** on HuR-bound transcripts during LPS treatment (8%). In absolute values, both PCs showed similar variance than at the transcriptome level (Supplementary Figure S9A). DEGs distributions in the considered comparison were visualized with a Venn diagram (Figure 4A, Supplementary Figure S9B). Similarly, for the RNA sequencing data a strong effect of LPS can be observed in the heat-map of differentially enriched genes, in which the unsupervised clustering separated LPS-treated samples from control samples, with the subcluster of **TM7nox** samples separately organized within the larger LPS branch (Figure 4B). As DHTS displaces transcripts with shorter 3’UTR and with lower of U/AU-rich elements than average^31^, we evaluated these two parameters in the comparisons between **TM7nox** and LPS co-treatment versus LPS and LPS vs DMSO. We observed that HuR-bound enriched mRNAs contained a longer 3’UTRs compared to the down-regulated ones as well as those not differentially regulated (Figure 4C, left).

**Figure 4.**
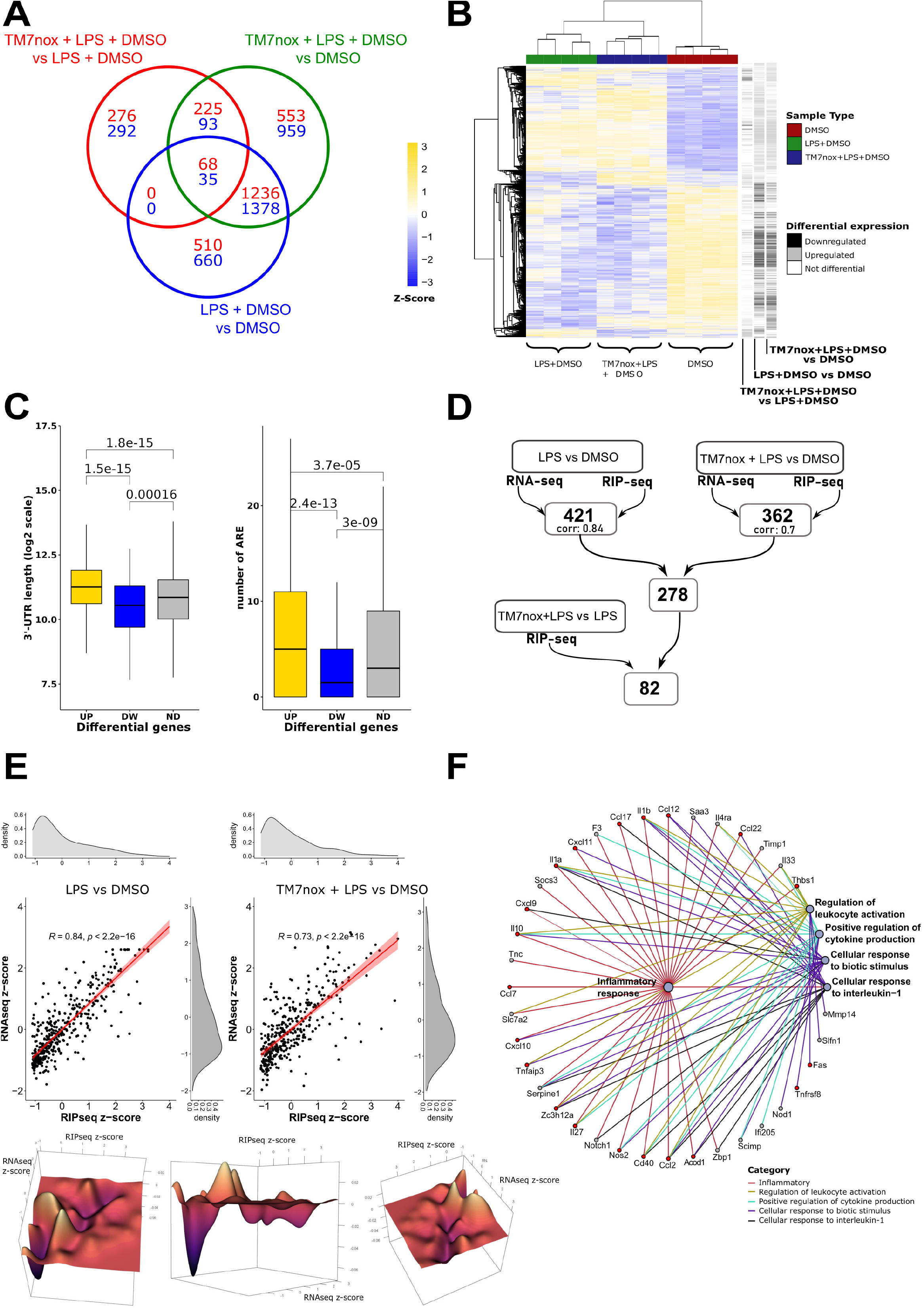
RIP-seq analysis reveals that TM7nox displaces the RNA targets from HuR. A) Venn of differentially expressed genes (DEGs), the numbers in each circle represent the number of DEGs between the different comparisons while the ones overlapping are for mutual DEGs (DMSO+LPS+**TM7nox** vs DMSO+LPS in green, DMSO+LPS+**TM7nox** vs DMSO in blue, DMSO+LPS vs DMSO in red). B) Heatmap of Z-score of differential genes across different samples, each grouping together with its own sample type (red: DMSO; green: DMSO + LPS; blue: DMSO + LPS + **TM7nox**). The track on the right shows differential gene expression between different comparisons (black: down-regulated DEGs; grey: up-regulated DEGs; white: no changes in DEGs). “Average” was used as clustering method and “correlation” for clustering distance of both rows and columns. C) (Left) Boxplot of 3’-UTR length of the up and down-regulated differentially expressed genes in the **TM7nox** and LPS co-treatment versus LPS and DMSO RIP-seq jointly with non-differential genes (ND). Up-regulated genes are in gold, down-regulated genes are in blue, non-differential genes are in gray. Wilcoxon test was performed between the three classes. (Right) Boxplot of the number of AREs of the up and down-regulated differentially expressed genes in the **TM7nox** and LPS co-treatment versus LPS and DMSO RIP-seq jointly with non-differential genes (ND). Up-regulated genes are in gold, down-regulated genes are in blue, non-differential genes are in gray. Wilcoxon test was performed between the three classes. D) Workflow of the genes filtering process starting from the LPS vs DMSO and **TM7nox** + LPS vs DMSO comparisons in both RNA-sequencing (*p-*adjusted < 0.05 & log_2_FoldChange > 1) and RIP-sequencing (*p-*adjusted < 0.05 & log_2_FoldChange > 3), identifying two different subsets of genes (421 genes with correlation value of 0.84 for LPS vs DMSO and 362 genes with correlation value of 0.73 for **TM7nox** + LPS vs DMSO comparison). 278 genes were thus identified to be commonly shared between these two subsets. Moreover, resulting from the filtering with the **TM7nox** + LPS vs LPS + DMSO down-regulated RIPseq comparison (*p-*adjusted < 0.05), 82 genes were found to be comprised of 20 genes of particular interest. E) (Left) Correlation between the RNA-sequencing z-score values and RIP-sequencing z-score values for the LPS and DMSO comparison. Each black dot represents a gene present in both experiments with a log2 FC > 1 for RNA-seq and log2 FC > 3 for RIP-seq. Regression line is depicted in red. Density plots of the distributions for each subset of values is shown on the right for RNA-seq z-score values and on the top for the RIP-seq z-score values. (Right) Correlation between the RNA-sequencing z-score values and RIP-sequencing z-score values for the **TM7nox** and LPS co-treatment versus DMSO comparison. Each black dot represents a gene present in both experiments with a log2 FC > 1 for RNA-seq and log2 FC > 3 for RIP-seq. Regression line is depicted in red. Density plots of the distributions for each subset of values is shown on the right for RNA-seq z-score values and on the top for the RIP-seq z-score values. (Bottom) 3D-rendered visualization from different angles of the difference between the correlation densities for LPS and DMSO comparison and TM7nox and LPS co-treatment versus DMSO comparison. On the axis, RNA-sequencing z-score, RIP-sequencing z-score values and distribution densities difference are shown. Color varies from purple to yellow according to the difference of the distributions values. F) Network visualization of the top five ranking GO pathways of subset of 82 genes of interest. Each pathway is represented by a gray dot and highlighted with a peculiar color, each gene is connected to the pathways it belongs to. Subset of 20 genes of interest is highlighted in red.

Moreover, the number of U/AU-rich regions present in HuR-bound mRNAs is higher compared to those which lost association or did not change (Figure 4C, right, Supplementary Table 4). Therefore, **TM7nox** displaces transcripts from HuR in a similar manner as DHTS. To get more insights on the mechanism of action of TMs, we investigated the correlation between induced gene transcription and HuR association (Figure 4D, E). 421 HuR-bound transcripts (log2 FC > 3) were also upregulated by LPS at the transcriptional level (Log2 FC > 1). We observed a strong correlation (coefficient = 0.84, Figure 4E left) between these two gene ensembles, corroborating the hypothesis of a HuR-mediated LPS response. GO analyses of these genes mirrored the LPS response at the transcriptome level, highlighting categories related to the innate immunity response, cytokine activity and chemotaxis. Many relevant genes mediating these responses were found associated to HuR, such as chemokines (*Cxcl2, Cxl10*), interleukins (*Il1a, Il1b, Il6, Il10*) and key regulators of immunity such as the Cd40 antigen, Janus Kinase2, and nitric oxide synthase 2.

By applying the same filtering process to **TM7nox** / LPS co-treated versus DMSO-treated samples, we found 362 upregulated genes, but we observed a decrease in the correlation coefficient to 0.73 (Figure 4E, right), suggesting an uncoupling effect of **TM7nox** on the association of each mRNA to HuR and its expression level. The difference between the two correlation densities was calculated and rendered in 3D using Delaunay triangulation and Dirchlet tessellation (Figure 4E, bottom). More in details, 278 out of the 362 genes were also present among the 421 transcripts of the previous LPS-treated dataset, but, despite an increase of the expression level and of HuR association compared to the DMSO / untreated level, 82 among these genes showed a decreased association to HuR in the **TM7nox** / LPS cotreatment condition with respect to the LPS condition. Interestingly, 47 genes out of 82 contain less AU rich motif than the average, suggesting that the decreased association to HuR is depending on the presence of the AU rich motif (Supplementary Table 1). GO analysis pointed to the inflammatory response as over-represented category (Figure 4F, Supplementary Table 3). Such genes are coherently modulated with a similar trajectory by **TM7nox** / LPS co-treatment, with *Cxcl10, Il1b, Cd40, Fas, Nos2* (Supplementary Figure S9D) as notable examples. As the correlation coefficient between expression and mRNA-HuR association decreased from 0.84 to 0.73 in the presence of **TM7nox**, we can infer that the latter modulates the ability of HuR to bind specific mRNAs. Therefore, this set of genes can be considered a core set of LPS-induced genes whose association to HuR has been dampened by **TM7nox**.

### TMs affect mRNA expression of LPS inducible genes and decrease binding of HuR to target RNAs

Thus, considering our sequencing results, we investigated the effect of three TMs to generalize the activity of TMs, as a co-treatment in response to LPS in murine RAW 264.7 macrophages at the single gene level. We co-treated cells with the active quinone species **TM6n, TM7nox** (10 µM), **DHTS** (5 µM) and LPS (1 µg/mL), and checked the expression levels of *Cxcl10, Cd40, Fas, Nos2* and *Il1b* by qRT-PCR at 6 hours post-treatment (Figure 5A). While LPS treatment induced the activation of the expression of several cytokines, **TM6n, TM7nox** and **DHTS** decreased their mRNA levels, which appeared to be significantly downregulated at 6 hours post co-treatments. Notably, *Cxcxl10, Cd40, Fas* and *Il1b* were significantly decreased, with *Nos2* showing a trend by treatment with **TM7nox** in HuR-IP samples, suggesting that TMs are indeed able to reduce the HuR-mRNA interaction within the cell (Figure 5B). To confirm the putative disruption of selected HuR-RNA complexes by TMs within cells, we performed RNA pull-down experiments using RAW 264.7 cells pre-treated for 6 hours focusing on **TM7nox** and DMSO as a control. Cell lysates were incubated with a biotinylated probe containing a 3’ ARE sequence belonging to *TNFα* 3’UTR, and a negative ARE-Neg biotinylated control containing a sequence that is not recognized by HuR. After incubation for 2 hours at 4°C, biotinylated probes were precipitated with streptavidin beads and HuR levels were assessed by Western Blot analysis (Figure 5C). Indeed, the HuR band in the **TM7nox**-treated sample precipitated with the *TNFα* ARE sequence was significantly decreased (∼ 50%) compared to both cells treated with DMSO and lysates precipitated with a HuR-unreactive ARE probe. This suggests that **TM7nox** competes with target RNAs for HuR binding within cells. BMDMs were co-treated with LPS (1 µg/mL) and **TM6n** or **TM7nox** (10 µM) for 6 hours. TMs treatment almost blocked LPS induced mRNAs expression, showing higher efficacy in BMDMs compared to RAW cells (Figure 5D). **TM7nox, TM6n** and control **DHTS** also decreased the release of Cxcl10 protein from RAW cells after 6 hours of treatment and post-LPS induction, as measured by ELISA (Figure 5E). Interestingly, **TM7nred** showed equivalent efficacy of **TM7nox** at the protein level in RAW cells (Figure 5E) and decreased Cxcl10 secretion in BMDMs (Figure 5F). Taken together, these results validate the RNA-seq and RIP-seq data and show the ability of TMs to interfere with LPS induced, HuR-mediated gene expression.

**Figure 5.**
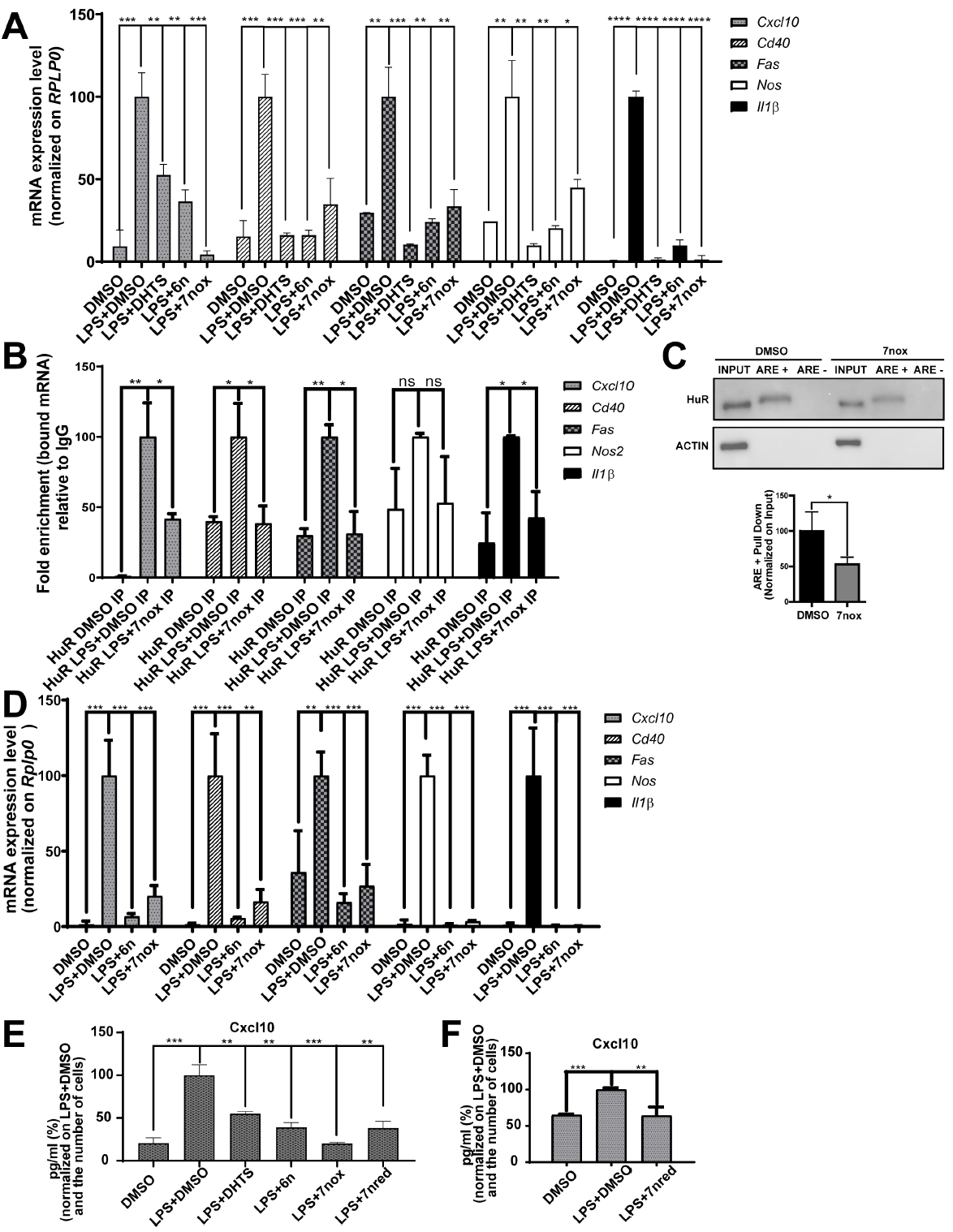
TM-7nox/7nred decrease the binding between HuR and identified targets, reducing their expression level and Cxcl10 secretion in both RAW 264.7 and BMDMs. A) RAW cells were co-treated for 6 hours with DHTS (5 μM) **TM6n** and **TM7nox** (10 μM), LPS (1μg/mL) and DMSO as control. mRNA levels of *Cxcl10, Cd40, Fas, Nos2 and Il1β* have been assessed through qRT-PCR using *RPLP0* as housekeeping gene and data have been normalized on LPS+DMSO condition. Data gave been plotted as mean ± SD of a biological quadruplicate (*p<0.05, **p<0.01, ***p<0.001, ****p<0.0001). B) TM7nox impairs the binding between HuR and identified targets. RIP-seq results have been validated by RNA Immunoprecipitation assay (RIP) followed by real time PCR (qRT-PCR). RAW 264.7cells were treated for 6 hours with DMSO alone and LPS (1μg/ul) +DMSO as control and LPS (1μg/mL) +**TM7nox** (10 μM). Subsequently, cells were lysed, and RNA precipitated with HuR antibody (IP) and IgG isotype (IgG) as negative control. Changes in the mRNAs bound to HuR in the control or treatment were evaluated through qRT-PCR and normalized on the corresponding values obtained with IgG, as negative control. The obtained numbers indicate the Fold enrichment; experiments were performed in a biological triplicate (*p<0.05, **p<0.01, ns= not significant). C) Pull Down assays performed in RAW 264.7 cells pre-treated with TM7nox or DMSO as control for 6 hours. Cell lysates were incubated for 1 hour at 4°C with either biotinylated probe containing HuR consensus sequence, either probe not supposed to bound by HuR as a negative control. Precipitations of the probes have been carried out with streptavidin beads and HuR level have been detected by WB analysis. HuR signal have been quantify on the Input (10%) and normalized on DMSO sample. Data have been plotted as mean ± SD of three independent experiments *p<0.05. D) BMDMs were co-treated for 6 hours with **TM6n** and **7nox** at 10 μM doses, LPS (1μg/mL) and DMSO as control. mRNA levels of *Cxcl10, Cd40, Fas, Nos2 and Il1β* have been assessed through qRT-PCR using *RPLP0* as housekeeping gene and data have been normalized on LPS+DMSO condition. Data gave been plotted as mean ± SD of a biological quadruplicate (*p<0.05, **p<0.01, ***p<0.001, ****p<0.0001). E) **TM7nox, TM6n**, DHTS and **TM7nred** treatment reduce Cxcl10 secretion in RAW 264.7 cell supernatants. Cxcl10 protein levels was measured with ELISA, RAW 264.7 cells were treated for 6 hours with DMSO as control, LPS (1μg/mL) plus DMSO or **TM7nred** (10 μM). Relative quantity of Cxcl10 pg/mL for each sample has been measuredon the number of cells quantified through crystal violet assay. After, data have been normalized on LPS+DMSO as control and number are expressed in percentage. Finally, data represent as mean ± SD of a biological triplicate (**p<0.01, ***p<0.001). F) TM7nred treatment reduces Cxcl10 secretion in BMDMs supernatants. Cxcl10 protein levels was measured with ELISA, RAW 264.7 cells were treated for 6 hours with DMSO as control, LPS (1μg/mL) plus DMSO or **TM7nred** (10 μM). Relative CXCL10 pg/mL for each sample have been measured on the number of cells quantified through crystal violet assay. After, data have been normalized on LPS+DMSO as control and number are expressed in percentage. Finally, data represent as mean ± SD of a biological quadruplicate (**p<0.01, ***p<0.001).

### TMs partially mimic HuR silencing and do not modulate NF-kB translocation

To investigate whether the activity of TMs in cells is due to an inhibition of HuR, we compared the effect of **TM7nred** and **TM7nox** with HuR silencing by 48 hours before LPS stimulation (6 hours) in RAW cells. In this case, we used both **TM7n** derivatives to show their biological equivalence in redox equilibration-compatible experimental protocols. Both compounds reduced the intracellular level of Cxcl10 and Il1b proteins, as measured by ELISA (Figure 6A). LPS did not induce the expression of HuR mRNA, while TMs reduced the expression of HuR mRNA both in DMSO/vehicle and LPS stimulation. This effect may be ascribed to the autoregulatory mechanism of HuR on its own mRNA. HuR silencing (Supplementary Figure S10A) in DMSO or LPS condition reduced the expression of HuR mRNA to about 50%, and both TMs did not show a significantly additive effect. The decrease of HuR did not reduce the expression level of *Cxl10, Cd40, Fas, Nos2* and *Il1b* mRNAs in basal conditions. However, during LPS stimulation *Cxcl10, Nos2* and *Il1b* expression levels were significantly decreased, while *Cd40* and *Fas* levels were not. Both TMs recapitulated the effect of HuR silencing during LPS stimulation for *Cxcl10, Nos2* and *Il1b*, but also decreased *Cd40* and *Fas* expression (Figure 6B). The simultaneous treatment with either **TM7n** derivative and HuR silencing did not show additive effect to HuR silencing alone on the protein level of Cxcl10 and Il1b (Supplementary Figure S10B). We then evaluated the stability of *Cxl10, Cd40, Fas, Nos2* and *Il1b* using two different protocols. In the first protocol, Actinomycin D (ActD) was co-administered with TM7nox 3 hours after LPS treatment; we evaluated the effect of **TM7nox** on the stability of the mRNAs irrespectively of its transcriptional impact. Contrary to expectations, comparing the expression levels of target RNAs at 1.5 hours after ActD we observed a trend of increased stabilization (Supplementary figure S10C). In a second experimental protocol, we administered **TM7nox** with LPS and added ActD after 3 hours. By comparing the expression levels of target RNAs at 1.5 and 3 hours after ActD we observed less mRNA in co-treated samples, suggesting a transcriptional impact of **TM7nox** during LPS co-treatment. Therefore, the effect of TM treatment resembles, but does not completely overlay with HuR silencing and the **TM7n** redox couple shows overlapping biological properties. While a 3 hours’ LPS treatment induces HuR into the cytoplasm^87^, co-treatment with our TMs did not counteract LPS-induced HuR shuttling. Similarly, **TM6n** and **TM7nox** (10 µM) did not induce HuR shuttling as previously observed in MCF-7 cells^30^ and did not counteract ActD-induced massive shuttling of HuR (Figure 7A). **DHTS**, as well, did not modulate HuR localization. The LPS induced NF-kb response was dampened by **TM7nox**, as suggested by functional analysis. To rule out that TMs directly inhibit the TLR4/NF-kB activity, we investigated whether NF-kB translocation into the nucleus was blocked by TMs, as this is the prerequisite for biological activity. Immunofluorescence experiments in RAW 264.7 cells (Supplementary Figure S10C, D), treated with **TM6n** or **TM7nox** for 3 hours, alone or in combination with LPS (1 µg/mL), indicated that LPS stimulates NF-kB nuclear localization, while TMs did not show any significant counteracting or stimulating action on its translocation (Figure 7B). DHTS partially reduced LPS induced nuclear shuttling of NF-kB^89^. Collectively, these data imply that TMs act independently from changing NF-kB cellular localization induced by LPS and are unable to stimulate HuR rescue in the nucleus as once inferred by drugs like ActD, suggesting once again that our TMs most likely function by modulating HuR-RNA binding activity.

**Figure 6.**
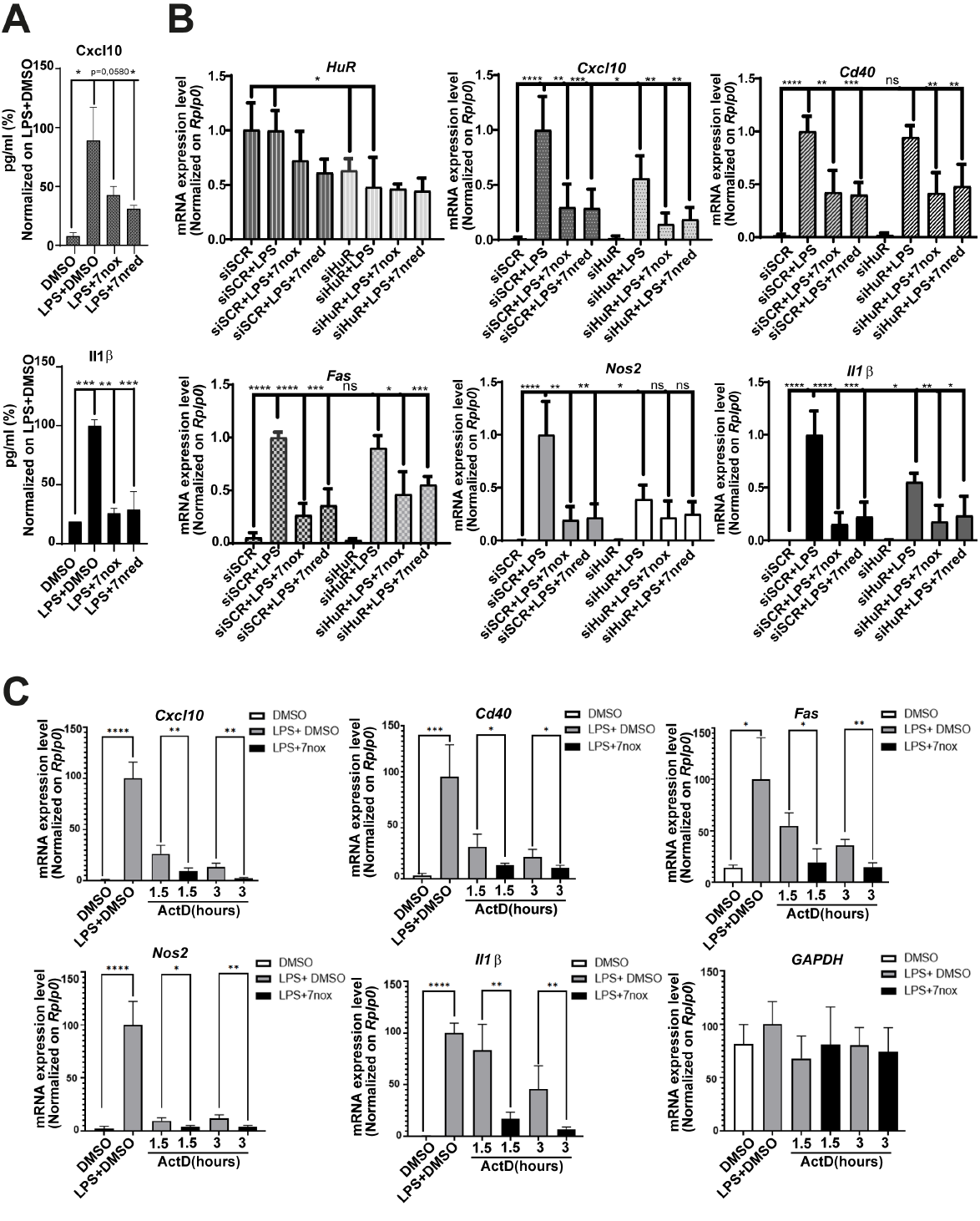
TM-7nox/7nred recapitulate partially HuR silencing without changing NFkB translocation and HuR subcellular localization. A) **TM7nox** and **TM7nred** treatment reduce Cxcl10 and Il1β intracellular levels in RAW 264.7cells. Protein levels were measured with ELISA, RAW 264.7 cells were treated for 6 hours with DMSO as control, LPS (1μg/mL) plus DMSO or TMs (10 μM). Respectively, 30 and 5 µg of cellular lysates were loaded to measure Cxcl10 and Il1β pg/mL. After, data have been normalized on LPS+DMSO as control and number are expressed in percentage. Finally, data represent as mean ± SD of a biological triplicate (**p<0.05, **p<0.01, ***p<0.001). B) qRT-PCR of target mRNAs in siSCR and siHuR, after 6 hours of co-treatment with LPS 1μg/mL plus DMSO, DMSO alone to control LPS stimulation, **TM7nox** and **7nred** 10 μM in RAW cells. Data have been plotted as Mean, and SD obtained are from 3 independent experiments (*p ≤0.05, **p ≤0.01, ***p ≤0.001, ****p<0.0001; ns, not significant). C) **TM7nox** affects transcription of LPS-induced cytokines. RAW cells were co-treated with with DMSO, LPS+DMSO, LPS+**TM7nox** for three hours. Act-D (2.5 μM) was then added/administered for other 1.5 or 3 hours. After, qRT-PCRs were performed to quantify the remaining *Cxcl10, Il1*β, *Cd40, Fas, Nos2* and *Gapdh* mRNA levels. Data are represented as mean ± SD of a biological triplicate (*p ≤0.05, **p ≤0.01, ***p ≤0.001, ****p<0.0001; ns, not significant).

**Figure 7.**
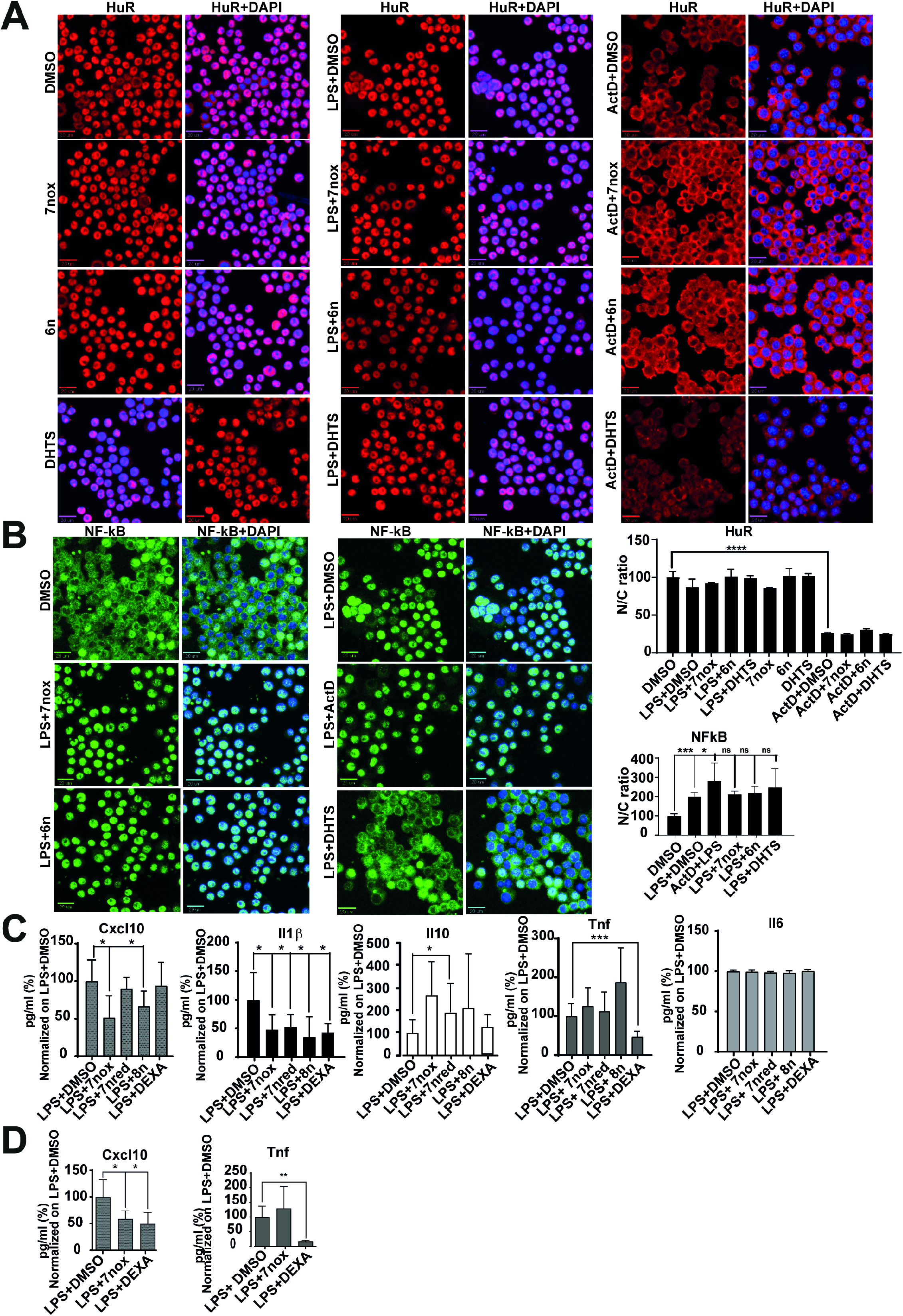
TMs modulate CXCL10 and Il1b secretion in LPS induced peritonitis mice models. A) Panel on the left show representative immunofluorescence HuR localization after LPS administration, single compounds treatments and in combination with LPS (1μg/mL). The right panel entails a representative immunofluorescence showing that HuR cytoplasm accumulation induced by Act-D is not changing upon treatments with different TMs (10 μM). Cells have been treated for 3 hours in combination with Act-D (2.5 μM). DMSO alone or in combination with Act-D and LPS has been used as control. In the graphs, the ratio of HuR fluorescent signal between nucleus and cytoplasm (N/C) is plotted. For image acquisition (40× high NA objective), Operetta was used and evaluation was carried out by selecting 13 fields/well. The N/C ratio represents the mean ± SD of single cells for every well (***p< 0.0001). B) Representative immunofluorescence showing that N-kB nuclear translocation inside the cells induced by LPS is not changing upon treatments with different TMs (10 μM). Cells have been treated for 3 hours in combination with LPS (1μg/mL). To obtain a positive control given by a high induction of NFkB related with a massive shuttling in the nucleus, we treated cells with 2.5 μM Act-D, and DMSO (CTRL) either alone or with LPS was used as negative control. In the graph, the ratio of NFkB fluorescent N/C signal is plotted. For image acquisition (40× high NA objective), Operetta was used and evaluation was carried out by selecting 13 fields/well. The N/C ratio represents the mean ± SD of single cells for every well (*p < 0.05; ***p < 0.0001). C) Levels of identified cytokines through Luminex analysis in sera from C57BL/6j wild type mice after administration of LPS (150 μg/25 g of body weight) and TMs (40 mg/kg) or DMSO for 2 hours. Data normalized on LPS+DMSO and expressed mean and SD as percentage of sex balanced mice group in which n= 6-8 (*p<0.05, ***p<0.001. D) CXCL10 and TNF levels measured with ELISA assays in the sera from C57BL/6j wild type mice after administration of LPS (150 μg/25 g of body weight) with TMs (40 mg/kg) or DMSO for 2 hours. DMSO alone has been used as a control of LPS inflammatory response insurgence and LPS+DMSO has been considered being drug vehicle. Bar graphs depict mean values ± SD from 5 mice per group (*p<0.05, **p ≤0.01)

### TMs effectively counteract LPS induced inflammatory response *in vivo*

Finally, *in vivo* efficacy of TMs was evaluated in the LPS-induced peritonitis mouse model^90^. Groups of six eight-weeks-old mice were co-treated with a sub-lethal dose of LPS and **TM7red, TM7nox** or its prodrug **TM8n** (i.p., 40 mg/kg); the redox couple was tested to evaluate the putative impact of different physico-chemical properties on *in vivo* potency, while the putative prodrug **TM8n** represented a possibly alternative route to *in vivo* efficacy. After 2 hours, mice blood was collected through cardiac puncture, serum was purified from plasma and cytokines were detected either by ELISA or by Luminex technology. We detected a Cxcl10, Il1b, Il6 and Tnfα cytokines increase in the mice sera after LPS treatment, validating the efficacy of LPS in activating the inflammatory response. Cytokines were tested in batch by Luminex, and dexamethasone was used as a counteracting agent to LPS induced inflammation. During co-treatment of LPS with TMs, Cxcl10 and Il1b levels decreased in sera derived from treated mice compared to control condition (LPS+DMSO); the anti-inflammatory cytokine Il10 showed an increasing trend that, however, did not reach statistical significance, while Il6 and TNFα did not show altered levels in mice sera (Figure 7C). A Cxcl10 decrease was also confirmed by ELISA assays in different mice cohorts (Figure 7D). TMs showed similar efficacy *in vivo*, and, notably, this confirmed our *in vitro* results for the inactive acetate prodrug **TM8n** to be converted, likely via esterases, to the **TM7nred** / **TM7nox** redox couple *in vivo* (Figure 7A). Finally, we observed that TMs behaved differently from dexamethasone in modulating LPS-induced response, as they were unable to modulate TNFα levels, were equally effective in reducing Il1b but were more potent in reducing Cxcl10 and increasing Il10 showing *in vivo* efficacy in this model.

## Discussion

After rationally designing four prospective TMs (**TM6n**, the redox couple **TM7nox** / **TM7nred**, and putative prodrug diacetate **TM8n**) targeted towards stronger potency on HuR and better bioavailability, we synthesized and submitted them to a preliminary *in vitro*-efficacy and physico-chemical characterization. As to the former, we confirmed HuR targeting for **TM6n** and **TM7nox** in cell-free and cellular assays; we observed the lack of cell-free activity for diacetate **TM8n** which, conversely, showed cellular activity in line respectively with easy interconversion of the redox couple **TM7nox** / **TM7nred** and a putative esterase-activated prodrug mechanism; and computational studies confirmed good affinity for HuR for oxidized orto quinones **TM6n** and **TM7nox**, while both diphenolic **TM7nred** and diacetate **TM8n** were predicted to be inactive. As to analytical studies, we confirmed the interconversion of the redox couple **TM7nox** / **TM7nred**.

A poor solubility was earlier reported for TMs^30^ and other HuR inhibitors^31,91^; nevertheless, a careful evaluation of the overall efficacy-stability-bioavailability profiles of our new TMs prompted us to run *in vivo* experiments for the **TM7nox** / **TM7nred** redox couple, and putative prodrug diacetate **TM8n**. TMs showed similar efficacy in terms of biochemical activity and cell viability; we selected the orto quinone **TM7nox** as a preferred tool for a detailed mechanistic characterization in terms of putative therapeutic effects on inflammation, keeping in mind its cellular / *in vivo* presence when reduced **TM7nred** was used. Indeed, we showed the overlapping biological activity of the two species of the redox couple. We investigated the ability of **TM7nox** in modifying the innate immune response triggered by LPS in macrophages. The most relevant mediator of LPS stimuli is the transcriptional activity of the LPS/TRL4/NF-κB axis. However, TMs blunt, but not abolish, such LPS-induced response without inhibiting the primary transcriptional response induced by NF-κB, as witnessed by our genome wide data. In addition, other RBPs as TTP, TIAR and hnRNP K are involved in the post-transcriptional response to LPS in macrophages^86^. In BMDM, PAR-CLIP and RNASeq analysis of the response to LPS indicated that HuR and TTP bind to specific target genes, but also compete for a small subset among them containing the binding motifs for each protein^33,88^. Therefore, it is plausible that the sole inhibition of HuR does not completely abrogate the LPS induced response. Furthermore, our RIP-seq experiments identified the most stably bound transcripts more likely engaged for translation, but, contrary to UV-crosslinked methods, it did not provide information about transient, low affinity interactions between HuR and a specific transcript^92^.

The modulation by TMs results, at least in part, from their ability to interfere with several HuR-AU-enriched target mRNAs interactions. We observed a comprehensive remodulation of the transcripts bound by HuR as, after **TM7nox** treatment, those that enriched their binding to HuR contain longer 3’-UTR and a higher number of U/AU rich regions compared to the ones that lost or did not change their binding. This mechanism of action is similar to what observed using DHTS in a different cellular model^31^. TMs’ treatment indeed decreases HuR binding to mRNAs of major players of the immune response such as *Cxcl10* and *Il1b*, leading to decreased secretion of the encoded proteins. CXCL10, known as interferon γ-induced protein 10 (IP-10), is strongly induced by IFNs (-α, -β but mostly -γ)^93,94^ as well as by LPS/NF-κB axis^95,96^, and other immune stimulants in monocytes, endothelial cells, fibroblasts, and cancer cells^97^. By interacting with the chemokine receptor CXCR3, CXCL10 stimulates differentiation of naive T cells to T helper 1 (Th1) cells and induces migration of immune cells to site of inflammation^98^. According to this, the inhibition of CXCL10 is considered beneficial in treating T cell mediated autoimmune diseases^93^ such as rheumatoid arthritis^99,100^ and type I diabetes (T1DM)^101^. On the same line, IL-1β belongs to the interleukin-1 family^102^ and is considered a potent pro-inflammatory cytokine mostly linked to innate immune response^103^. It is a clinical target in autoinflammatory diseases such as cryopyrin-associated periodic syndromes (CAPS) or autoimmune disease^104^. Our data have been collected by co-administering LPS with **TM7nox**. In so doing, as highlighted in the ActD chase experiments, we observed that TMs modulate the transcription of a core set of LPS target genes by interfering with HuR nuclear activity. The same experiments where TMs are administered after LPS stimulus, suggest also a post-transcriptional mechanism that, for the genes investigated, leads to the stabilization of the genes of interest. Further studies are required to investigate this aspect. These limitations notwithstanding, our observations suggest the putative usefulness of TMs in preclinical models of autoinflammatory and autoimmune diseases. A similar effect has been reported by the small molecule HuR modulators DHTS^31^ and SRI-42127^85^. **TM7nox** disrupts the interaction between HuR and its target mRNAs similarly to DHTS, and the latter was shown to exhibit a strong anti-inflammatory activity by suppressing mRNA expression or secreted protein levels of the pro-inflammatory mediators TNFα and IL6 in LPS-induced RAW and BMDMs cells, as well as *in vivo* in LPS-induced mouse models^105–108^. The anti-inflammatory activity of DHTS has also been associated with its ability to suppress nuclear translocation of NF-κB^109^, that was not observed in the case of TMs, suggesting a partially different mechanism of action between the two classes of small molecules. In LPS-activated primary microglial cells, SRI-42127 reduces the expression and release of several cytokines and chemokines such as IL-1β, IFN-γ, CXCL1, CCL3, IL-6 but an opposite effect on CXCL10^85^. Notably, the spectrum of affected target genes is largely overlapping with the ones affected by TMs, although the mechanism of HuR interaction is different for the two chemotypes. In fact, SRI-42127 is an inhibitor of HuR homodimerization and a blocker of HuR nucleo-cytoplasmic shuttling, that abolishes the cytoplasmic function of HuR, reducing the stability of targeted mRNAs^85^; conversely, TMs, and in particular orto quinone **TM7nox** presented here, bind to HuR between the first two RRMs, competing with target mRNAs and locking the protein in a closed conformation, with no apparent influence on dimerization and nucleo-cytoplasmic shuttling. As shown by our molecular dynamic simulations, **TM7nox** not only binds between the RMM1-2 domains, but also induces the surrounding loops of the RNA binding cavity to fold around the ligand to dramatically reduce the available buried surface area between the two RRM domains. Thus, such conformational state surely hampers the accommodation of the binding mRNA.

TMs, as shown by RNA-seq and RIP-seq experiments, change the HuR-bound transcriptome and modulate the activity of HuR. As functional data reported here suggest that TMs have a detectable effect while a strong HuR activation takes place, such as after LPS exposure, their putative usefulness during different stress stimuli that lead to HuR activation should be investigated.

## Acknowledgments

The authors wish to thank the LaBSSAH - CIBIO Next Generation Sequencing Facility of the University of Trento for sequencing samples and HTS facility for immunofluorescence analyses.

